# Neurons in Visual Cortex are Driven by Feedback Projections when their Feedforward Sensory Input is Missing

**DOI:** 10.1101/2020.01.24.919142

**Authors:** Andreas J Keller, Morgane M Roth, Massimo Scanziani

## Abstract

We sense our environment through pathways linking sensory organs to the brain. In the visual system, these feedforward pathways define the classical feedforward receptive field (ffRF), the area in space where visual stimuli excite a neuron^1^. The visual system also uses visual context, the visual scene surrounding a stimulus, to predict the content of the stimulus^2^, and accordingly, neurons have been found that are excited by stimuli outside their ffRF^3–8^. The mechanisms generating excitation to stimuli outside the ffRF are, however, unclear. Here we show that feedback projections onto excitatory neurons in mouse primary visual cortex (V1) generate a second receptive field driven by stimuli outside the ffRF. Stimulating this feedback receptive field (fbRF) elicits slow and delayed responses compared to ffRF stimulation. These responses are preferentially reduced by anesthesia and, importantly, by silencing higher visual areas (HVAs). Feedback inputs from HVAs have scattered receptive fields relative to their putative V1 targets enabling the generation of the fbRF. Neurons with fbRFs are located in cortical layers receiving strong feedback projections and are absent in the main input layer, consistent with a laminar processing hierarchy. The fbRF and the ffRF are mutually antagonistic since large, uniform stimuli, covering both, suppress responses. While somatostatin-expressing inhibitory neurons are driven by these large stimuli, parvalbumin and vasoactive-intestinal-peptide-expressing inhibitory neurons have antagonistic fbRF and ffRF, similar to excitatory neurons. Therefore, feedback projections may enable neurons to use context to predict information missing from the ffRF and to report differences in stimulus features across visual space, regardless if excitation occurs inside or outside the ffRF. We have identified a fbRF which, by complementing the ffRF, may contribute to predictive processing.

---

To characterize the classical feedforward receptive field (ffRF), we first mapped receptive field locations of layer 2/3 excitatory neurons in primary visual cortex (V1) in awake head-fixed mice with two-photon calcium imaging (Fig. 1a). To determine the center of a neuron’s ffRF, we used circular patches of drifting gratings presented individually at different locations (Fig. 1b; see Methods). To estimate the size of the neuron’s ffRF, we obtained a size tuning function by varying the diameter of the grating (from 5 to 85°; Fig. 1c) centered on the neuron’s ffRF and presented at the neuron’s preferred orientation (see Methods). The responses were maximal for gratings of 13.1 ± 0.4° in diameter (mean ± SEM; 1190 neurons in 9 mice) and were strongly suppressed with increasing grating size (90 ± 1% suppression at 85°; mean ± SEM; 1190 neurons in 9 mice). Thus, consistent with previous reports, layer 2/3 excitatory neurons in V1 had a ffRF diameter greater than 10° on average^9–12^ and their responses were suppressed when a stimulus extended beyond the ffRF to cover surrounding regions^13–15^.

**Figure 1.**
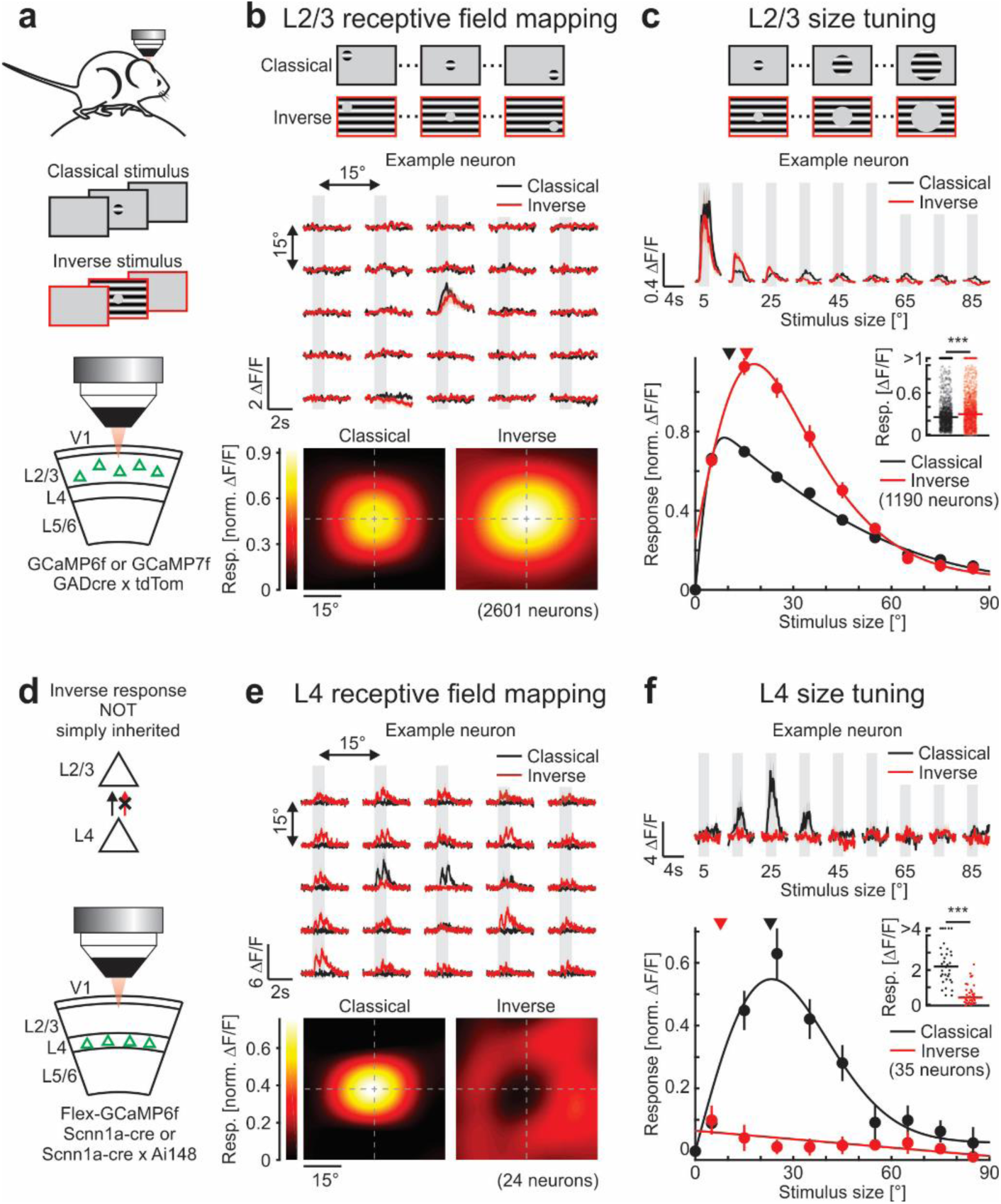
Layer-specific responses to inverse stimuli. **a**, Experimental configuration. Classical or inverse stimuli are presented to awake mice while imaging calcium responses in layer 2/3 excitatory neurons of primary visual cortex (V1). **b**, Top: Schematic of receptive field mapping. Center: Trial-averaged calcium responses from an example neuron for each stimulus location. Here and in all other figures, black and red traces are responses to classical and inverse stimuli, respectively, and shaded areas are periods of stimulus presentation. Bottom: Population-averaged receptive field for responses to classical or inverse stimuli aligned to the center of the classical feedforward receptive field (ffRF). Note the strong overlap of the receptive fields mapped with classical or inverse stimuli (2601 neurons in 9 mice). **c**, Top: Schematic of stimuli used for size tuning functions. Center: Trial-averaged calcium responses from an example neuron for each stimulus size. Classical and inverse stimuli are centered on the ffRF. Bottom: Size tuning functions to classical and inverse stimuli. For every neuron, both size tuning functions are normalized by their maximum response to classical stimuli. Solid lines are fits to the data (see Methods). Triangles indicate median preferred size. Inset: Maximum responses. Horizontal lines, medians. Wilcoxon signed-rank test; *: p < 10^-4^; 1190 neurons in 9 mice. **d**, Schematic of results and experimental configuration. **e** and **f**, As above but for layer 4 excitatory neurons (24 neurons in 4 mice). Note that, in contrast to (**b**), the responses to inverse stimuli surrounds the ffRF (**e**). Also note the monotonically decreasing inverse size tuning function in (**f**) as compared to (**c**). Wilcoxon signed-rank test; ***: p < 10^-6^; 35 neurons in 6 mice. Traces and data points represent mean ± SEM (shading or error bars, respectively). Here and in all other figures, error bars are present but sometimes smaller than symbols.

To determine the spatial extent of the suppressive regions, we presented a full-field grating in which a portion was masked by a gray circular patch (Fig. 1a, b). We reasoned that if a large grating suppresses the excitatory neuron’s response because it stimulates suppressive regions surrounding the ffRF, the response of the excitatory neuron should be partially recovered when a gray patch is placed on a suppressive region, i.e. when part of the suppressive region is not stimulated. We varied the location of the gray patch along the same grid used to determine the center of the neuron’s ffRF (see Methods). By averaging the responses to these stimulus grids across neurons, we obtained two separate activity maps; one of the ffRF and one of the suppressive regions. Unexpectedly, the peak of the map of the suppressive regions overlapped with the peak of the ffRF map (Fig. 1b). This suggests that, at the population level, the largest recovery from suppression was obtained when the gray patch was located at the center of the ffRF.

Given the relatively low resolution of our maps, the overlap between the two maps could have arisen by averaging individual neurons each with a suppressive zone at a different location yet adjacent to the ffRF. To obtain a finer measure of the response amplitude of a neuron to the gray patch, we placed the patch on the center of the neuron’s ffRF and systematically varied its diameter. Even the smallest tested size of the gray patch (5°) placed on the center of the ffRF evoked a response that was larger than the full-field grating indicating that the resolution of the maps is unlikely to impact our results. Strikingly, neuronal responses increased with the size of the gray patch up to 16.6 ± 0.4° (mean ± SEM; 1190 neurons in 9 mice) and then progressively diminished with larger diameters (Fig. 1c). These responses were not a result of the sharp edges of the stimuli since we observed similar responses when blurring the edges of the border between the grating and the gray patch (Extended Data Fig. 1). Thus, the dependence of a neuron’s response to the size of a gray patch on a full-field grating was similar to that of a grating patch on a gray background.

These results show that layer 2/3 excitatory neurons are not only excited by classical stimuli (grating patches on a gray background) but are also robustly excited by inverse stimuli (full-field gratings with a gray patch) centered on a neuron’s ffRF. These two stimuli, the classical and the inverse, had an antagonistic impact on layer 2/3 neurons because the responses of layer 2/3 neurons to both stimuli together (e.g. a full-field grating) were significantly smaller than the responses to either of the stimuli presented alone (Wilcoxon signed-rank test; p < 10^-10^; for both classical and inverse stimuli of 15° compared to a large diameter grating, 85°; Fig. 1c). We thus defined a neuron as inverse tuned if its response to at least one inverse stimulus of any size centered on its ffRF was significantly larger than that to a full-field stimulus (see Methods). According to this criterion, 79% of the visually responsive excitatory neurons in layer 2/3 were inverse tuned (943 of 1190 neurons in 9 mice; Fig. 1a-c and Extended Data Fig. 2). In addition, we computed the inverse tuning index (ITI; 0: classical only; 0.5: equal peak responses to classical and inverse stimuli; 1: inverse only; see Methods). Layer 2/3 excitatory neurons had a unimodal ITI distribution peaking at 0.52 ± 0.01 (mean ± SEM; Extended Data Fig. 3a).

In inverse-tuned neurons, there is a region surrounding the ffRF that, when stimulated in the presence of a stimulus in the ffRF, is suppressive while, when stimulated in the absence of a stimulus in the ffRF, is excitatory. We first compared the tuning properties of the surrounding excitatory region with those of the ffRF in layer 2/3 neurons using classical and inverse stimuli at different orientations and directions (Extended Data Fig. 3b). On average, the orientation tuning to inverse stimuli was sharper than the one to classical stimuli, and individual neurons showed a wide range of orientation and direction selectivity indexes for classical and inverse stimuli (Extended Data Fig. 3b-f). Moreover, individual neurons were not necessarily tuned to the same orientation when stimulated by classical or inverse stimuli (Extended Data Fig. 3g), indicating that the surrounding excitatory region has its own set of tuning properties. We determined the interaction between the surrounding excitatory region and the ffRF in neurons with similar orientation preference by varying the contrast of classical or inverse stimuli presented simultaneously (Extended Data Fig. 3h). With matching contrasts above 13% the mean activity in layer 2/3 excitatory neurons was lower than when presenting either classical or inverse stimuli alone, indicating an overall antagonistic interaction between the surrounding excitatory area and the ffRF.

Is inverse tuning in layer 2/3 excitatory neurons inherited from earlier stages of cortical processing? To this end, we imaged the activity of excitatory neurons in layer 4, the main cortical feedforward input layer, by conditionally expressing GCaMP6f (Fig. 1d-f). As for layer 2/3 excitatory neurons, we first estimated the ffRF center and the size dependence of the response for each neuron using classical stimuli. We then used inverse stimuli to determine the location of the suppressive regions. In striking contrast to what we observed in layer 2/3, the suppressive regions of layer 4 neurons surrounded their ffRF, creating a ring around the center (Fig. 1e). Further, the responses of layer 4 neurons to inverse stimuli placed on the center of the ffRF decreased monotonically with the size of the stimulus, which was markedly different from layer 2/3 neurons (Fig. 1c vs Fig. 1f). This decrease in the response of layer 4 neurons is consistent with the progressive reduction of feedforward drive (Extended Fig. 2). Thus, we found no evidence of inverse tuning in layer 4 neurons (ITI: 0.10 ± 0.02; mean ± SEM). Overall, the spatial organization of suppressive regions of layer 4 neurons relative to their ffRFs is in agreement with previous models and observations^7, 16^ and importantly, distinct from layer 2/3 neurons.

Could our inability to resolve a suppressive ring around the ffRF of layer 2/3 neurons be due to an insufficient spatial resolution of the mapping stimuli (20° patches with a 15° spacing)? We compared our receptive field estimates to those obtained with a finer grid (10° patches with a 5° spacing) and found that both estimations were similar (Extended Data Fig. 4a, b). Additionally, we specifically analyzed layer 2/3 neurons with larger ffRFs, comparable to the sizes observed in layer 4 (i.e. preferred size >15°; Extended Data Fig. 4c, d). The classical size tuning function of this subpopulation was almost indistinguishable from that of layer 4 neurons (Extended Data Fig. 4d). Yet, in contrast to layer 4 neurons, their suppressive region still colocalized with the peak of the ffRF (Extended Data Fig. 4c). Thus, inverse tuning in layer 2/3 excitatory neurons is not simply inherited from layer 4 neurons and hence represents an operation that emerges along the cortical laminar hierarchy.

Sources of input to layer 2/3 neurons are neuron-type specific^14, 17^. Is inverse tuning also present in layer 2/3 inhibitory neurons? To determine whether the three major classes of cortical inhibitory neurons, the parvalbumin- (PV), vasoactive-intestinal-peptide- (VIP) and somatostatin-expressing (SOM) neurons, are also inverse tuned, we conditionally expressed GCaMP6f in each of these three classes and characterized their responses to classical and inverse stimuli (Fig. 2). For both PV and VIP cells the classical and the inverse maps peaked and overlapped around the center of the ffRF (Fig. 2b, c). Furthermore, both PV and VIP cells showed surround suppression to classical stimuli, consistent with previous reports^14^ and size tuning functions to inverse stimuli that peaked well above the response of the neuron to the largest classical stimuli. In contrast, SOM cells only showed, on average, a poor and spatially diffuse response to inverse stimuli (Fig. 2d), little surround suppression to classical stimuli^14^ and no response to inverse stimuli above the response to the largest classical stimuli. Thus, the majority of PV and VIP neurons were inverse tuned (97%; 58 of 60; ITI: 0.52 ± 0.02; mean ± SEM and 58%; 43 of 74; ITI: 0.42 ± 0.02; mean ± SEM, respectively), similar to layer 2/3 excitatory neurons, while most SOM were not (30% inverse tuned; 54 of 179; ITI: 0.22 ± 0.02; mean ± SEM).

**Figure 2.**
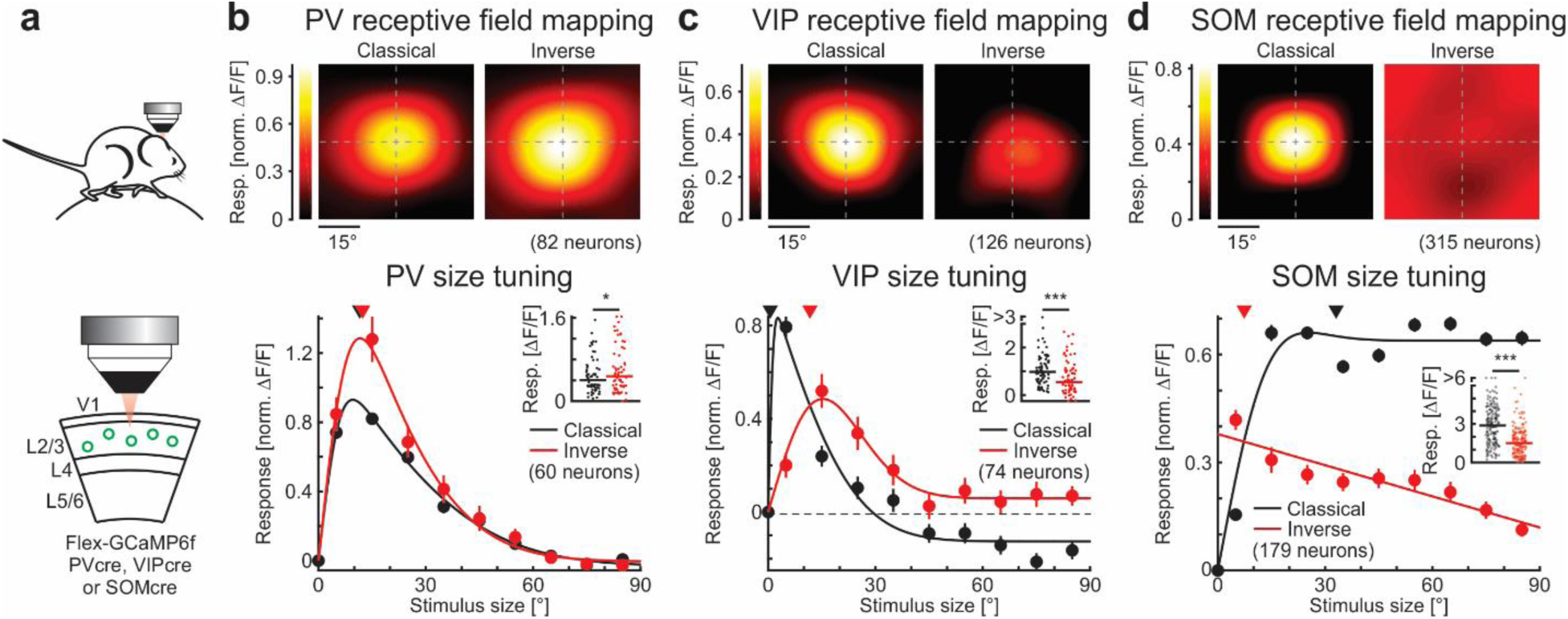
Neuron-type specific response to inverse stimuli. **a**, Experimental configuration. Classical or inverse stimuli are presented to awake mice while imaging calcium responses in layer 2/3 parvalbumin-positive (PV), vasoactive-intestinal-peptide-positive (VIP), or somatostatin-positive (SOM) inhibitory neurons in V1. **b**, Top: Population-averaged receptive field for PV inhibitory neuron responses to classical or inverse stimuli aligned to the center of the ffRF (82 neurons in 6 mice). Bottom: Size tuning functions to classical and inverse stimuli. For every neuron, both size tuning functions are normalized by their maximum response to classical stimuli. Solid lines are fits to the data (see Methods). Triangles indicate median preferred size. Inset: Maximum responses. Horizontal lines, medians. Wilcoxon signed-rank test; *: p = 0.021; 60 neurons in 7 mice. **c**, Same as in (**b**) but for VIP inhibitory neurons. Top: 126 neurons in 4 mice. Bottom: Wilcoxon signed-rank test; ***: p = 3.6 × 10^-4^; 74 neurons in 8 mice. **d**, Same as in (**b**) but for SOM inhibitory neurons. Top: 315 neurons in 5 mice. Bottom: Wilcoxon signed-rank test; ***: p < 10^-10^; 179 neurons in 5 mice.

Inverse tuning is therefore a ubiquitous property of layer 2/3 neurons but absent in layer 4. Even though our results show that inverse-tuned layer 2/3 neurons do not directly inherit this property from layer 4, the excitatory region surrounding these layer 2/3 neurons could still be generated via feedforward inputs from layer 4 with spatially offset ffRFs (Fig. 3a). Alternatively, the excitatory region surrounding layer 2/3 neurons could be generated via feedback projections, as feedforward projections only represent the minority of excitatory inputs onto those neurons^18, 19^. To address this, we compared the latency of the responses of inverse-tuned neurons to classical and inverse stimuli using extracellular electrophysiological recordings. We inserted a multichannel linear probe into V1 (Fig. 3a) and used classical stimuli to determine the ffRF location of the recorded neurons, as described for calcium imaging. This approach allowed us to record isolated single units throughout the cortical layers, including infragranular layers (layer 5/6). Interestingly, we found that a large fraction of infragranular units (50%, 60 of 119 units; ITI: 0.40 ± 0.03; mean ± SEM) were also inverse tuned (Extended Data Fig. 5). In inverse-tuned units recorded in both supra- and infragranular layers the time course of the responses to classical and inverse stimuli were markedly different, as shown by their peristimulus time histograms (PSTHs; Fig. 3b). While the time course of the response to classical stimuli was characterized by a fast initial transient followed by a response plateau, the response to inverse stimuli showed a significantly slower progression towards steady state (Fig. 3c-e, see Methods). Further, the response to inverse stimuli was delayed relative to the classical response (50 ± 20 ms; mean ± SEM; 15 units; Fig. 3f). The same biases were observed when comparing the response dynamics to classical stimuli in all responsive neurons to those to inverse stimuli in inverse-tuned neurons (Fig. 3g). This difference in latencies and the slower dynamics of responses to inverse stimuli suggest that inverse tuning may involve additional processing steps and is thus unlikely to emerge from the feedforward pathway.

**Figure 3.**
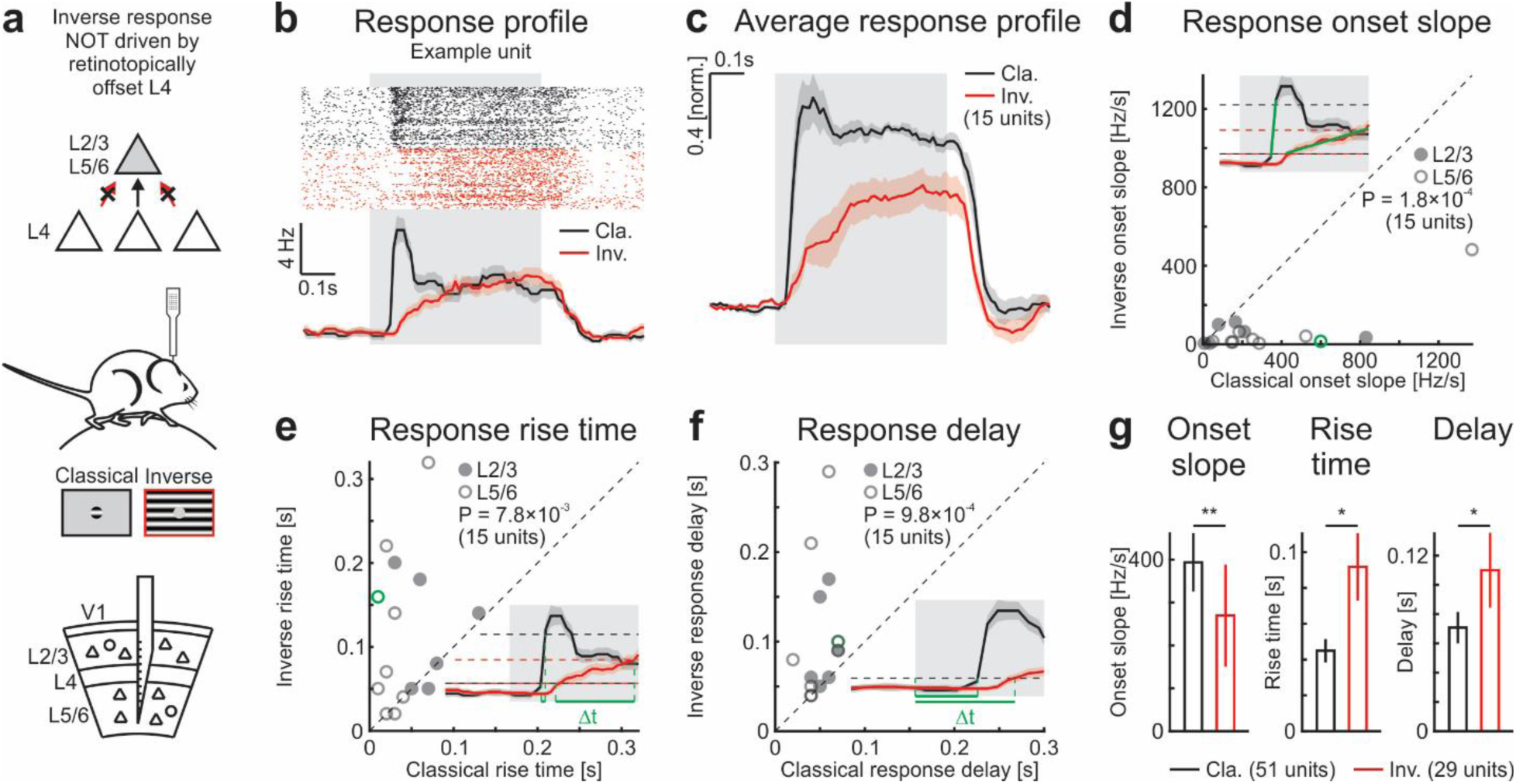
Slow and delayed responses to inverse stimuli. **a**, Schematic of results and experimental configuration for extracellular recordings in V1. **b**, Response of an example unit in layer 5/6 to classical and inverse stimuli centered on the ffRF. Top: Raster plot for classical and inverse stimuli (1000 trials each). The initial phase of the drifting grating was randomized for each trial. Bottom: Peristimulus time histogram (PSTH; 10 ms bins). **c**, Population average of normalized PSTHs (15 units in 4 mice). For every unit, PSTH was normalized by the average activity to the classical stimulus. **d**, Scatter plot of the onset slope of the response to classical and inverse stimuli. Closed and open symbols are units from layers 2/3 and 5/6, respectively. Green symbol represents the example unit shown in (**b**). Inset: PSTH from (**b**) to illustrate differences in slopes. Horizontal dotted lines are lower and upper thresholds to compute the slopes (see Methods). Darker symbols are overlapping data points. Wilcoxon signed-rank test; p = 1.8 × 10^-4^; 15 units 4 mice. **e**, Same as (**d**) but for rise time. Wilcoxon signed-rank test; p = 7.8 × 10^-3^; 15 units in 4 mice. **f**, Same as (**d**) but for delay. Wilcoxon signed-rank test; p = 9.8 × 10^-4^; 15 units in 4 mice. **g**, Mean onset slopes, rise times, and delays for classical and inverse stimuli of units that were tuned to either the classical or the inverse stimulus (see Methods). Wilcoxon rank-sum test; onset slope, **: p = 1.1 × 10^-3^; rise time, *: p = 0.017; delay, *: p = 0.031; classical: 51 units in 8 mice; inverse: 29 units in 8 mice. Traces and bars represent mean ± SEM (shading or error bars, respectively).

Could the responses to inverse stimuli depend on feedback projections from higher visual areas (HVAs)? To this end, we harnessed the impact of anesthesia on the sensory responses of neurons, which has been suggested to be more pronounced in HVAs than in V1^20^. We verified the differential effect of anesthesia by comparing the impact of isoflurane on the responses to classical stimuli in V1 and in HVAs identified beforehand using wide-field intrinsic imaging (see Methods). Indeed, responses were significantly more suppressed by anesthesia in HVAs than in V1 (Extended Data Fig. 6). If the responses of layer 2/3 neurons to inverse stimuli rely on feedback projections, they should therefore be more sensitive to anesthesia than the responses of the same neurons to classical stimuli. We imaged layer 2/3 excitatory neurons first in awake mice while presenting classical and inverse stimuli, and then imaged the same neurons under isoflurane anesthesia (Fig. 4a). In inverse-tuned neurons, anesthesia strongly suppressed the responses to inverse stimuli (peak response reduction of 64 ± 4%; mean ± SEM; 49 neurons; Fig. 4b, c) while the responses to classical stimuli were only weakly affected (peak response reduction of 7 ± 13%; mean ± SEM; same 49 neurons; Fig. 4b, c). This was also the case for neurons whose peak response was larger to inverse than to classical stimuli (Fig. 4c). Thus, anesthesia has a stronger impact on the amplitude of the responses to inverse stimuli, consistent with the hypothesis that it is mainly driven by feedback projections.

**Figure 4.**
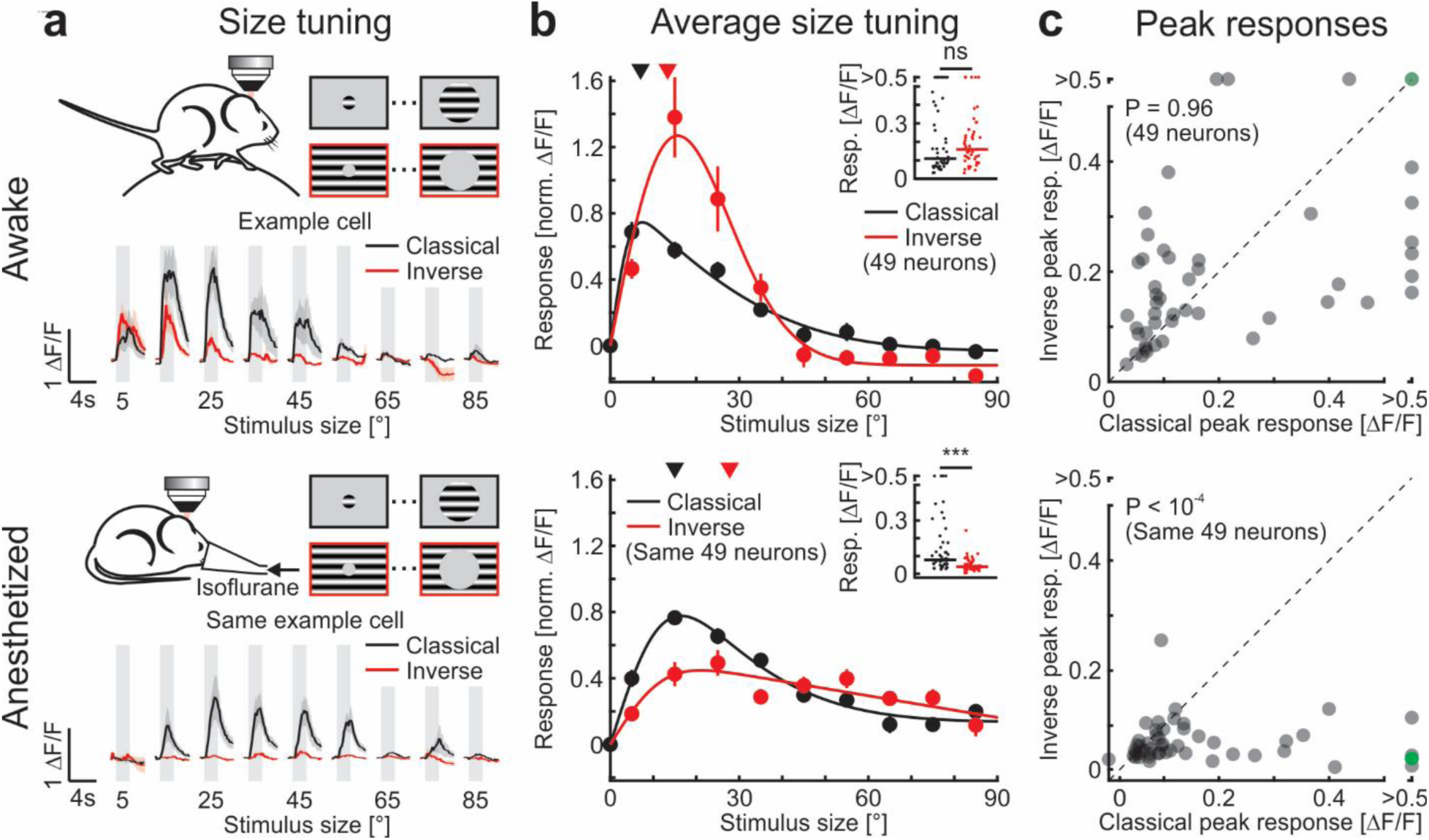
Anesthesia preferentially reduces responses to inverse stimuli. **a**, Top diagram: Experimental configuration as in Fig. 1a. Top traces: Calcium responses of an example neuron to classical and inverse stimuli of different sizes in an awake mouse. Bottom: Same example neuron but under isoflurane anesthesia. Note the larger reduction of responses to inverse than to classical stimuli. **b**, Population-averaged size tuning functions for classical and inverse stimuli in awake (top) and anesthetized (bottom) mice (top and bottom: same 49 neurons). For every neuron, each size tuning function is normalized by the maximum awake response to the classical stimulus. Solid lines are fits to the data (see Methods). Triangles are median preferred size. Insets: Maximum responses. Horizontal lines, medians. Top: Wilcoxon signed-rank test; ns: p = 0.96; 49 neurons in 5 mice. Bottom: Wilcoxon signed-rank test; ***: p < 10^-4^; same 49 neurons. **c**, Scatter plot of peak responses of inverse-tuned neurons to classical and inverse stimuli in awake (top) and anesthetized (bottom) mice (top and bottom: same 49 neurons). Green symbol represents the example neuron shown in (**a**). Wilcoxon signed-rank test; top: p = 0.96; bottom: p < 10^-4^; 49 neurons in 5 mice. Traces and data points represent mean ± SEM (shading or error bars, respectively).

To directly test the involvement of top-down feedback projections on inverse tuning, we optogenetically silenced HVAs while recording extracellular electrophysiological activity in V1 (Fig. 5a-e). We silenced cortical activity by scanning a laser beam across HVAs to optogenetically activate inhibitory neurons expressing Channelrhodopsin-2 (ChR2; Fig. 5a). Control recordings confirmed that this procedure efficiently silenced cortical activity in areas targeted by laser stimulation (Extended Data Fig. 7a-c). In inverse-tuned units, silencing HVAs reduced spontaneous activity and responses to classical stimuli, in particular those of smaller diameters and, as previously shown, increased the activity for large stimuli in surround suppressed units^21–23^ (Fig. 5b-d; Extended Data Fig. 8). The response to inverse stimuli, however, was strongly suppressed (Fig. 5b-e). Upon silencing HVAs, inverse stimuli evoked responses reminiscent of those observed in layer 4 that decreased with increasing size of the gray patch, as would be expected by the progressive masking of a stimulus. These effects cannot be explained by a direct impact of scattered laser light on V1 as the effect of laser stimulation on ChR2-expressing inhibitory neurons decreased rapidly with distance. Indeed, the stimulation was insufficient to increase the activity of ChR2-expressing inhibitory units in V1 beyond 400 μm from the center of stimulation or when silencing HVAs (Extended Data Fig. 7d-g; see Methods). Silencing HVAs by activating local inhibitory neurons, however, could also impact V1 activity if these inhibitory neurons were to send long-range projections to V1. To test this possibility, we injected a retrogradely transported virus to conditionally express a fluorescent reporter in inhibitory neurons projecting to the injection site (Extended Data Fig. 7h-l). Consistent with previous reports^24^, we observed very few retrogradely labelled inhibitory neuron somata in HVAs (HVAs: 4.3 ± 0.9 neurons; V1: 510 ± 10 neurons; mean ± SEM; 3 mice) indicating that direct inhibitory projections from HVAs to V1 are rare and therefore unlikely to impact our results.

**Figure 5.**
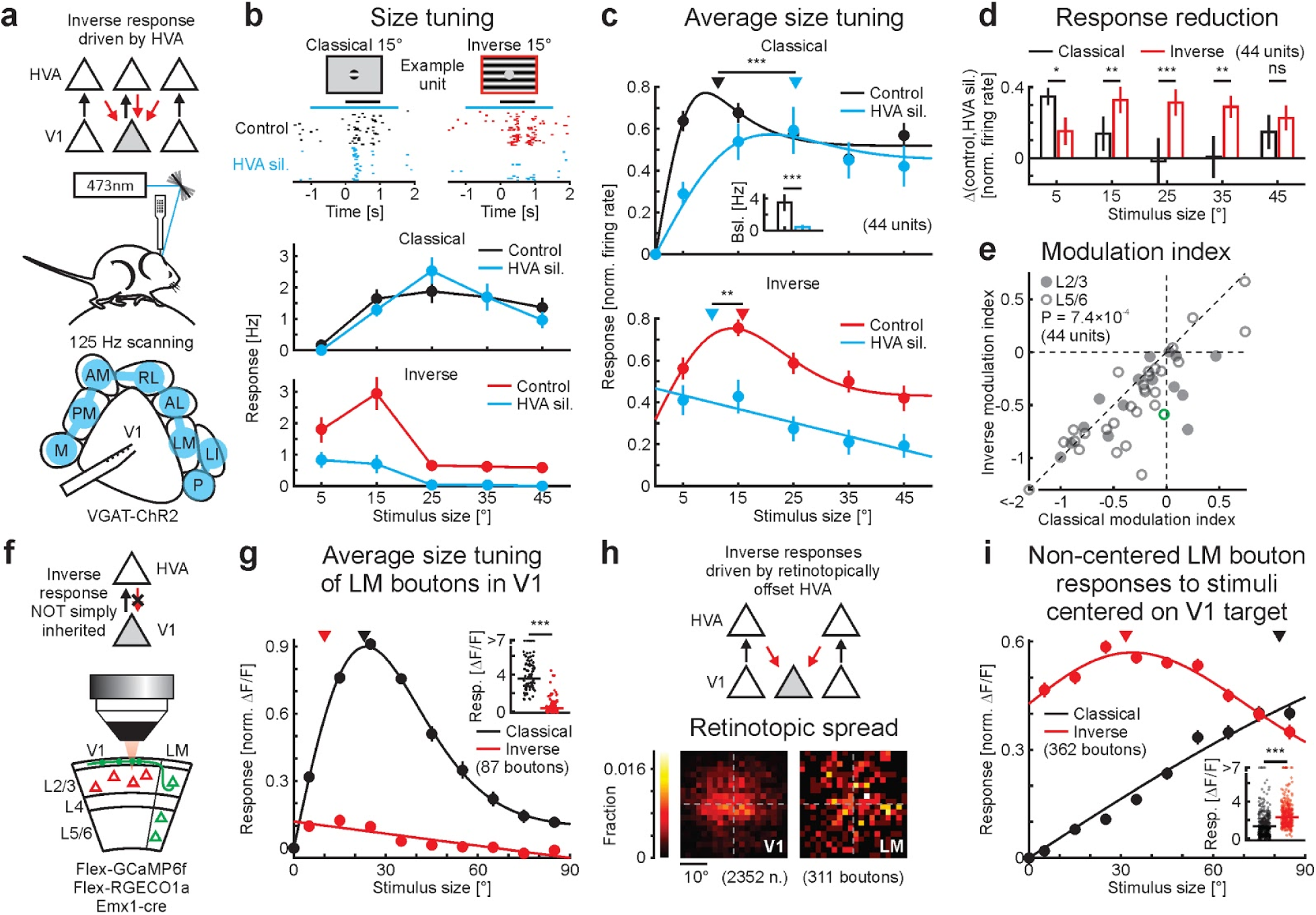
Higher visual areas contribute to inverse tuning in V1. **a**, Schematic of results and experimental configuration. A laser beam is scanned over higher visual areas (HVAs) around V1 for optogenetic silencing while recording in V1. **b**, Top: Raster plot of an example unit in layer 5/6 in response to classical and inverse stimuli of 15° under control conditions and during silencing of HVAs (HVA sil.; blue; 30 trials each). Black and blue horizontal lines are periods of stimulus presentation and HVA silencing, respectively. Bottom: Size tuning function of an example unit (baseline subtracted firing rates) to classical and inverse stimuli and during silencing of HVAs (blue). Note the stronger reduction in response to inverse stimuli upon silencing HVAs. **c**, Population-averaged size tuning function to classical (top) and inverse (bottom) stimuli in control and during optogenetic silencing of HVAs (blue). For every unit, baseline subtracted size tuning functions are normalized by their maximum control response to classical (top) or inverse stimuli (bottom). Solid lines are fits to the data (see Methods). Triangles indicate median preferred size for each condition. Wilcoxon signed-rank test; classical, ***: p = 3.2 × 10^-4^; inverse, **: p = 3.0 × 10^-3^; 44 units in 12 mice. Inset: Mean baseline firing rate under control conditions and during silencing of HVAs (blue). Wilcoxon rank-sum test; ***: p < 10^-9^; 44 units in 12 mice. **d**, Difference in firing rates (baseline subtracted and normalized) between control conditions and HVA silencing for classical and inverse stimuli for the different sizes tested. Wilcoxon signed-rank test; 5°, *: p = 8.0 × 10^-3^; 15°, **: p = 1.4 × 10^-3^; 25°, ***: p < 10^-4^; 35°, **: p = 3.5 × 10^-3^; 45°, ns: p = 0.70; 44 units in 12 mice. **e**, Scatter plot of the modulation indexes of HVAs silencing for responses to classical and inverse stimuli (see Methods). Closed and open symbols are units from layer 2/3 and 5/6, respectively. Green symbol represents the example neuron shown in (**b**). Wilcoxon signed-rank test; p = 7.4 × 10^-4^; 44 units in 12 mice. **f**, Schematic of results and experimental configuration. **g**, Population-averaged size tuning function for classical and inverse stimuli of LM boutons that are retinotopically aligned with their V1 target (see Methods). Note that these LM boutons only weakly respond to inverse stimuli. Solid lines are fits to the data (see Methods). Triangles are median preferred size. Insets: Maximum responses. Horizontal lines, medians. Wilcoxon signed-rank test; ***: p < 10^-10^; 87 boutons in 5 mice. **h**, Top: Schematic of results. Bottom: Retinotopic spread of the ffRF of V1 neurons and LM boutons (2352 neurons and 311 boutons in the same 5 mice). **i**, Population-averaged size tuning function of LM boutons (375 boutons in 5 mice) that are NOT retinotopically aligned with their V1 target and have at least one significant response to any inverse stimuli. Note that both classical and inverse stimuli were presented at the ffRF location of their putative V1 targets (see Methods) and NOT at the ffRF location of the imaged LM boutons. Solid lines are fits to the data (see Methods). Triangles are median preferred size. Insets: Maximum responses. Horizontal lines, medians. Data points and bars represent mean ± SEM.

Do HVAs contribute equally to inverse responses in V1? To test this, we silenced individual HVAs while recording single unit responses in V1 to classical and inverse stimuli (Extended Data Fig. 9). While the silencing of several visual areas reduced the response to inverse stimuli, the strongest stimulus-specific effect on inverse responses was observed when silencing the lateromedial visual area (LM).

Is inverse tuning directly inherited from inverse-tuned neurons in HVAs? To address this question, we expressed GCaMP6f in LM and RGECO1a in V1 to determine the responses properties of LM axonal boutons in layer 1 of V1 while mapping the retinotopic coordinates of the V1 site (Fig. 5f-i; Extended Data Fig. 10a-h). LM boutons whose receptive fields were centered on the retinotopic coordinates of the imaged V1 site (11% of all visually responsive boutons; 87 of 800), showed surround suppression to classical stimuli and were not inverse tuned (ITI: 0.14 ± 0.01; mean ± SEM; Fig. 5g). When presenting an inverse stimulus centered on the receptive field of the LM boutons, the response decreased with increasing diameter of the gray patch, similar to what observed in layer 4 neurons in V1 (Fig. 1f). This was not a general property of LM neurons since directly imaging cell bodies in LM showed inverse tuning in some of the neurons, and in the population average (ITI: 0.33 ± 0.02; mean ± SEM; Extended Data Fig. 10i-k). Thus, V1 neurons do not directly inherit inverse tuning from LM because those LM neurons that project back onto matching retinotopic coordinates in V1 show no inverse tuning.

Inverse tuning of layer 2/3 neurons could result from LM inputs which, while *per se* not inverse tuned, have spatially offset receptive fields relative to those of layer 2/3 neurons. These LM inputs would respond to an inverse stimulus centered on the V1 retinotopic coordinates because their receptive field, being offset relative to the gray patch, would be directly stimulated by the grating. We mapped the spatial offset of the receptive field of LM boutons relative to the retinotopic coordinates of the V1 sites. The receptive field centers of LM boutons showed a wide scatter relative to the retinotopic coordinates of the V1 site, larger than the scatter of receptive field centers of layer 2/3 neurons at the V1 site, and consistent with a previous study^25^ (Fig. 5h; Extended Data Fig. 10f, g). Thus, LM boutons with spatially offset receptive field centers converge on a given retinotopic site in V1. If these LM inputs contribute to the inverse response of layer 2/3 neurons, they should respond to inverse stimuli centered on the V1 site. This was indeed the case on average and a large fraction of these boutons significantly responded to inverse stimuli (51% of the non-aligned boutons; 362 of 711; or 45% of all visually responsive boutons; 362 of 800; Extended Data Fig. 10h). Furthermore, these boutons responded robustly to all tested inverse stimuli, predominantly to small stimulus sizes (Fig. 5i), consistent with the inverse size tuning in layer 2/3 neurons (Fig. 1c). In addition, the response of these boutons to progressively larger stimuli centered on the V1 site increased gradually (Fig. 5i), consistent with their receptive fields being offset relative to the center of the stimulus. Thus, these results show that inverse tuning in layer 2/3 V1 neurons likely results from the feedback of non-inverse-tuned neurons in HVAs whose receptive fields are offset relative to the receptive field centers of the V1 neurons they converge on.

In conclusion, our results demonstrate that feedback projections to V1 neurons generate a second, distinct excitatory receptive field that surrounds the ffRF. This feedback receptive field (fbRF) is absent in layer 4, the main feedforward input layer, and emerges along the laminar processing hierarchy in the supra- and infragranular layers of V1. Neurons in V1 receive feedback projections originating from HVAs, yet whether these projections merely modulate or are actually capable of driving V1 neurons has remained a matter of debate^15, 26, 27^. Our data clearly show that feedback from HVAs is capable of driving V1 neurons and that this drive comes, at least in part, from retinotopically offset neurons. Further, we show that the fbRF and the ffRF are mutually antagonistic when stimulated with uniform stimuli and that this antagonism is not exclusive to excitatory neurons but also present in inhibitory neurons expressing PV and VIP. Through this antagonism, neurons respond when a stimulus is presented in either the fbRF or ffRF but not in both together, effectively performing an exclusive-OR operation. Suppression of responses to stimuli in the ffRF by surrounding stimuli is a well-established phenomenon that enables neurons to report boundaries by detecting differences in stimulus features between the excited region inside the ffRF and their surround^7, 8, 14, 15, 28–31^. Our results show that neurons with an excitatory fbRF report differences in stimulus features regardless of whether the excited region is located inside or outside the ffRF. This may represent a generalization in the ability to detect boundaries across visual space. While the mechanisms underlying the mutual antagonism between the fbRF and the ffRF remains to be elucidated, we hypothesize that SOM inhibitory neurons, which, in contrast to PV and VIP inhibitory neurons, respond poorly to inverse stimuli while robustly responding to large stimuli covering both the fbRF and the ffRF, could be responsible for this operation. Indeed, functional elimination of SOM neurons has been shown to relieve excitatory neurons from the suppression mediated by large stimuli^14^.

Our data establish a role of HVAs in the generation of the fbRF, however, we cannot exclude a contribution of local excitation within V1^32, 33^. Regardless, the presence of a fbRF may underlie phenomena such as filling-in or illusory contours in which the stimulus in the ffRF is absent or obstructed^4, 5, 34–36^. Similarly, a fbRF may account for the reported interactions between excitatory regions outside and inside the ffRF as a basis for contextual modulation^8, 15, 28, 37, 38^, detection of borders^6, 7^ or pop-out effects^39^.

The antagonism between feedback excitation descending from HVAs and feedforward excitation ascending from the periphery is reminiscent of models of predictive processing^27, 40, 41^. In these models, bottom-up information about the stimulus is compared with top-down predictions based on experience, such that only errors between prediction and stimulus identity are represented and passed along to update the prediction. Surround suppression has been interpreted within the framework of predictive processing in space because if the visual stimulus surrounding the ffRF of a neuron is a good predictor for the stimulus in the ffRF there is no error between stimulus prediction and identity and, therefore, the response of that neuron is suppressed^40^. Without inverse tuning, as in layer 4 neurons, errors are only generated when there is a stronger stimulus present in the ffRF than expected based on the surround (e.g. no stimulus in the surround), but not for the inverse situation when the stimulus is weaker in the ffRF than expected based on the surround (e.g. stimulus only in the surround). With inverse tuning, as in layer 2/3 neurons, the framework of predictive processing generalizes to stimuli within and outside of the ffRF due to the presence of a fbRF. It will be important to determine the extent to which visual experience is necessary to generate the fbRFs.

## Acknowledgements

We thank M. Mukundan, B. Wong, and L. Bao for technical support, R. Beltramo for helping with extracellular recordings, J. Isaacson, G. Keller, R. Nicoll, and M. Heindorf for comments on the manuscript and the members of the Scanziani laboratory for helpful discussions of this project as well as for comments on the manuscript. This project was supported by the NIH grant U19NS107613, the Howard Hughes Medical Institute and the Swiss National Science Foundation grants P300PA_177882 and P2EZP3_162284 to A.J.K and P300PA_177898 to M.M.R. Confocal images were acquired at the Nikon Imaging Center at UCSF.

## Author contributions

A.J.K. and M.S. designed the study. A.J.K. and M.M.R. conducted all experiments and analysis. M.S., A.J.K. and M.M.R wrote the manuscript.

## Methods

### Animals

All experimental procedures were conducted in accordance with the regulation of the Institutional Animal Care and Use Committee of the University of California, San Francisco. Mice of either sex were kept on a C57BL/6 background (except VIP-IRES-cre) and were of the following genotype:

Gad2-IRES-cre (GAD2^tm2(cre)Zjh^ ; JAX:010802) × Ai14 (Gt(ROSA)26Sor^tm14(CAG-tdTomato)Hze^; JAX:007914) for imaging of layer 2/3 excitatory neurons (9 mice; Fig. 1a-c and 4, Extended Data Fig. 1, 3, and 4); Emx1-IRES-cre (Emx1^tm1(cre)Krj^ ; JAX:005628) for imaging layer 2/3 excitatory neurons and axons from LM (5 mice; Fig. 5f-i, and Extended Data Fig. 10a-h); Gad2-IRES-cre (GAD2^tm2(cre)Zjh^ ; JAX:010802) for imaging layer 2/3 neurons and labelling inhibitory projections (8 mice; Extended Data Fig. 6, 7h-k, and 10i-k); PV-cre (Pvalb^tm1(cre)Arbr^; JAX:017320) × Ai14 (Gt(ROSA)26Sor^tm14(CAG-tdTomato)Hze^; JAX:007914) for imaging of layer 2/3 parvalbumin-expressing inhibitory neurons (PV; 7 mice; Fig. 2b); VIP-IRES-cre (Vip^tm1(cre)Zjh^; JAX: 010908) × Ai14 (Gt(ROSA)26Sor^tm14(CAG-tdTomato)Hze^; JAX:007914) for imaging of layer 2/3 vasoactive-intestinal-peptide-expressing inhibitory neurons (VIP; 8 mice; Fig. 2c); Sst-IRES-cre (Sst^tm2.1(cre)Zjh^; JAX:028864) × Ai14 (Gt(ROSA)26Sor^tm14(CAG-tdTomato)Hze^; JAX:007914) for imaging of layer 2/3 somatostatin-expressing inhibitory neurons (SOM; 5 mice; Fig. 2d); Scnn1a-Tg3-cre (Tg(Scnn1a-cre)3Aibs/J; JAX:009613) and Scnn1a-Tg3-cre (Tg(Scnn1a-cre)3Aibs/J; JAX:009613) × Ai148 (Igs7^tm148.1(tetO-GCaMP6f,CAG-tTA2)Hze^; JAX:030328) for imaging layer 4 excitatory neurons (5 mice and 1 mouse, respectively; Fig. 1d-f, and Extended Data Fig. 4d); and VGAT-ChR2-EYFP (Tg(Slc32a1-COP4*H134R/EYFP)8Gfng/J; JAX:014548) for electrophysiology and optogenetic inhibition experiments (20 mice; Fig. 3 and 5a-e, Extended Data Fig. 5, 7a-g, 8, and 9). The mice were housed on a reverse light cycle (light/dark cycle: 12/12 hrs). At the start of the experiments, all mice were older than 2 months.

### Viruses

We injected the following viruses: AAV2/1.ef1a.GCaMP6f.WPRE (FMI Vector Core Facility), AAV2/1.ef1a.DIO.GCaMP6f.WPRE (FMI Vector Core Facility), AAV2/1.CAG.CGaMP6f (Janelia Vector Core), AAV2/9.syn.GCaMP7f (Addgene), AAV1.Syn.Flex.NES-jRGECO1a.WPRE.SV40 (Addgene) and AAVretro.CAG.Flex.tdTomato (Addgene). Viruses were diluted to use titers of approximately 5 × 10^12^ genome copies/ml and 50nl were injected at each injection site (3 to 5 for two-photon and 1 for anatomy experiments) and each depth (2 from 350 to 200 μm below the pial surface for two-photon calcium imaging experiments; 4 from 650 to 200 μm below the pial surface for the anatomy experiments and two-photon recordings of LM boutons).

### Surgery

Mice were anesthetized with 2% isoflurane or with a mixture of Fentanyl (West-Ward Pharmaceuticals, 0.05 mg/kg), Midazolam (Akorn, 5.0 mg/kg) and Dexmedetomidine (Zoetis, 0.5 mg/kg), injected subcutaneously. Mice’s body temperature was monitored and kept constant. To prevent the eyes from drying, a layer of lubricant ointment (Rugby) was applied. The skin above the skull was disinfected with povidone iodine. For mice prepared for intrinsic optical imaging (those needed for two-photon calcium imaging in higher visual areas (HVAs) or in LM boutons, and for all electrophysiology experiments), the bone over the right visual cortex was thinned, the exposed skull was covered with a thin layer of glue (krazy glue) and a headplate was attached using dental cement (Ortho-Jet Powder, Lang). The mice were then allowed to recover for several days before any other surgical or experimental procedures. For two-photon experiments, a craniotomy was made over the right visual cortex (3 to 4.5 mm in diameter) and viruses were injected with a micropump (UMP-3, World Precision Instruments) at a rate of 2 nl/s. The craniotomy was then sealed with a glass coverslip using cyanoacrylate glue and, if not already present, a headplate was attached. For electrophysiology experiments, a small craniotomy was performed (approximately 0.3 mm in diameter) guided by the activity maps of the visual cortex obtained by intrinsic optical imaging. After the recording, the mouse was either perfused for histology or its skull was protected with Kwik-Cast (World Precision Instruments) for the next experiment. For anatomical experiments, the skin was sutured after the viral injection using 6-0 suture silk (Fisher Scientific NC9134710). To reverse the anesthesia induced by the Fentanyl-Midazolam-Dexmedetomidine mixture, a mixture of Naloxone (Hospira, 1.2 mg/kg), Flumazenil (West-Ward Pharmaceuticals, 0.5 mg/kg), and Atipamezol (Zoetis, 2.5 mg/kg) was injected subcutaneously after the surgical procedures.

### Visual stimulation

Visual stimuli were generated using the open-source Psychophysics Toolbox based on Matlab (MathWorks). Stimuli were presented at a distance of 15 cm to the left eye on a gamma-corrected LED-backlit LCD monitor (DELL) with a mean luminance of 20 cd/m^2^. For two-photon experiments using a resonant scanner, the power source of the monitor’s LED backlight was synchronized to the resonant scanner turnaround points (when data were not acquired) to minimize light leak from the monitor^42^. We presented drifting sinusoidal gratings (2 Hz, 0.04 cycles/°, 100% contrast) unless stated otherwise. The trial structure of all stimulus sessions (receptive field mapping, size tuning, …) was block randomized (the block size was given by the total number of parameter combinations). In all raster plots (Fig. 3, Fig. 5, and Extended Data Fig. 7), we separated stimulus conditions for clarity.

#### Intrinsic imaging

To estimate the visual area locations and their retinotopic maps using intrinsic imaging, we presented a narrow white bar (5°) on a black background, slowly drifting (10°/s) in one of the cardinal directions (10 to 20 trials per direction). In addition, we presented 25° patches of gratings at different retinotopic locations (usually one nasal and one temporal, 20 trials each). Gratings were presented for 2 s at 8 different directions (0.25 s each) followed by 13 s of gray screen.

#### Receptive field mapping

Stimuli consisted of either a 20° circular grating patch on a gray screen (classical stimulus) or a 20° gray circular patch on a full-field grating (i.e. large gratings covering the entire screen, approximately 120 × 90°; inverse stimulus) with a 15° spacing between the center of the patches (regular grid). For two-photon calcium imaging experiments, stimuli were presented for 1 s at a single direction or for 2 s at the four cardinal directions (0.5 s each). Stimulation periods were interleaved by 2 s of gray screen. We recorded 5 to 10 trials per stimulus condition. For electrophysiological experiments, stimuli were presented for 0.5 s at a single direction interleaved by 1 s of gray screen. We recorded 20 trials per stimulus condition. In addition, we used a finer grid of grating patches in a subset of experiments (patches of 10° with a spacing of 5°; Extended Data Fig. 4a, b).

#### Orientation tuning

We presented gratings of at least 15° diameter drifting in 8 directions (5 to 10 trials and 20 trials per direction for two-photon calcium imaging and electrophysiology experiments, respectively). For Extended Data Fig. 3, we additionally presented inverse gratings drifting in 8 directions, centered on the classical feedforward receptive field (ffRF). Stimulus presentation time was 1 s interleaved with 1.5 to 2 s of gray screen.

#### Size tuning

Patches of gratings and inverse gratings were displayed at 9 different sizes, equally spaced from 5 to 85° in diameter (10 trials per size; for two-photon experiments) or at 5 different sizes, equally spaced from 5 to 45° in diameter (20 to 30 trials per size; for electrophysiology experiments), centered on the ffRF. Stimulation time was either 2 s interleaved by 4 s of gray screen (for two-photon experiments) or 1 s interleaved by 1.5 s of gray screen (for electrophysiology experiments). Trials with optogenetic stimulation had an additional 1 s pre-stimulus and 0.5 s post-stimulus gray screen during which the optogenetic light source was turned on and the total number of trials was doubled (**Optogenetics** below). In addition, we blurred the edge of the patches using a sigmoid function rising from 1% to 99% over 10° in a subset of experiments (Extended Data Fig. 1). All other parameters were the same as for the size tuning described above.

#### Contrast tuning

We simultaneously presented classical and inverse stimuli with several test contrasts (0, 2^-6^, 2^-5^, …, 1). Stimuli were presented for 2 s interleaved by 4 s of gray screen (10 trials per stimulus combination).

#### Response dynamics

To estimate the temporal response profile to inverse stimuli (Fig. 3), we presented patches of gratings and inverse gratings at a single size (1000 trials each). These gratings were presented either at 15° or 20°, for 0.5 s interleaved by 1 s of gray screen. The initial phase of the drifting gratings was randomized to avoid overestimating the onset delay of the response for simple-cell-like receptive fields.

### Behavioral monitoring

All mice were habituated (3 to 5 days) to experimental rigs before starting experiments. During all awake experiments, we recorded the positions of the left eye using a CMOS camera (DMK23UM021, Imaging Source) with a 50 mm lens (M5018-MP, Moritex), tracked the running speed of the mouse, and monitored its general behavior using a webcam (LifeCam Cinema 720p HD, Microsoft). Excluding eye-movement or running trials did not affect the results. For experiments under anesthesia that followed awake experiments (Fig. 4 and Extended Data Fig. 6), mice were anesthetized with isoflurane (approximately 1% in O_2_) delivered with a nose cone. After induction of anesthesia, mice’s body temperature was monitored and kept constant. To ensure an adequate depth of anesthesia, we tracked the left eye and monitored general behavior.

### Intrinsic optical imaging

We used intrinsic optical imaging to identify the center of primary visual cortex (V1) or the locations of HVAs. We sedated the mice with chlorprothixene (0.7 mg/kg) then lightly anesthetized with isoflurane (0.5 to 1% in O_2_) delivered through a nose cone. The rectal temperature was monitored and maintained at 37°C. We illuminated visual cortex with 625 nm light from two LED light sources (M625F2, Thorlabs) using 1.5 mm light fibers (FP1500URT, Thorlabs). The intrinsic optical signal was measured with an Olympus MVX stereo-macroscope using a narrow bandpass filter (700/13 nm BrightLine, Semrock). We acquired the images at 10 Hz with a CCD camera (Orca-Flash 4.0 v2, Hamamatsu) using custom-written software in LabVIEW (National Instruments).

### Two-photon calcium imaging

Imaging was performed using either a galvanometric-scanner based MOM (Sutter) or a resonant-scanner based (8 kHz) Bergamo II two-photon microscope (Thorlabs), both controlled by ScanImage (Vidrio). Using the MOM system, we acquired images of 128 × 128 pixels at a single depth at 5.92 Hz frame rate. With the Bergamo II, we acquired images of 380 × 512 pixels at 1 or 4 depths at 40 Hz or 8 Hz frame rate, respectively. We obtained similar results with both systems, so all data were pooled. The illumination light source was a Ti:sapphire laser (Chameleon Ultra II, Coherent) used at an excitation wavelength of 910 nm for green indicator imaging and of 1040 nm for red indicator imaging. The laser power under the objective (16×, Nikon) never exceeded 50 mW (laser pulse width 140 fs at a repetition rate of 80 MHz).

### Electrophysiology

We performed extracellular recordings using multi-electrode silicon probes (A1×32-Edge-5mm-20-177-A32, NeuroNexus) with 32 channels spaced by 20 μm. The recording electrodes were controlled with micromanipulators (Luigs&Neumann) and coated with DiO lipophilic dyes (Life Technologies) for *post hoc* identification of the electrode track. We recorded the bandpass-filtered (0.1 Hz to 7.5 kHz) signals at 30 kHz using an Intan system (RHD2000 USB Interface Board, Intan Technologies).

### Optogenetics

We used the VGAT-ChR2-EYFP mouse line to ensure a homogeneous expression of the opsin. To silence parts of visual cortex, we used a 473 nm laser (LuxX 473-80, Omicron-Laserage). The light was first guided through a pinhole to collimate the beam, then sent through a long-range focal lens (AC254-300-A, Thorlabs) to focus the light onto the cortical surface (theoretical spot size ≤ 200 μm), before it entered a 2D-galvo system (GVS202, Thorlabs) to direct the light to the regions of interest. The scanners were controlled by custom-written software in LabVIEW (National Instruments) and guided by a CMOS camera (DMK23UM021, Imaging Source) with a 50 mm lens (M5018-MP, Moritex). For Fig. 5a-e and Extended Data Fig. 7-9, we defined 8 HVAs (P, LI, LM, AL, RL, AM, PM, and M) based on the intrinsic optical imaging maps established before the optogenetic experiment (see **Data analysis**, *Intrinsic optical imaging maps*). These areas were consecutively scanned in a circular manner with a dwell time of ≤ 1 ms per area (resulting in 125 Hz frequency for the whole cycle). The laser was briefly shut off each time the beam moved from area M and P to avoid silencing parts of V1 (see **Visual Stimulation** for timing within a trial). For assessing the role of single HVAs in the generation of inverse tuning, we targeted each area individually (Extended Data Fig. 9). To verify the effectiveness of the silencing using this approach, we performed control recordings by scanning over the recording site in V1 (Extended Data Fig. 7a-c). To measure the spatial extent of silencing, we parked the laser at 5 locations at and around the recording site (800 μm and 400 μm lateral and medial of the recording site and on the recording site itself, randomizing which location to silence for each trial), targeted individual HVAs, or scanned over these 8 HVAs (Extended Data Fig. 7d-g). For experiments scanning over all 8 HVAs, the laser power was set to approximately 0.75 mW/mm^2^ (total power at the surface of cortex was 3 mW distributed over approximately 4 mm^2^ of illuminated HVAs). For experiments targeting individual locations or HVAs, the laser power was set to approximately 4 mW/mm^2^ (total power at the surface of cortex: 2 mW).

### Histology

Mice were deeply anesthetized with 5% isoflurane and urethane, and transcardially perfused with PBS followed by 4% paraformaldehyde in PBS. The brain was then embedded in 2 to 3% agar and 100 μm thick sections were cut using a microtome (Leica VT1000 S vibratome). Slices were mounted using a Vectashield HardSet mounting medium containing DAPI (H-1500-10, Vector Laboratories H1500). Images were acquired with an Olympus MVX10 MacroView microscope or a Nikon Ti CSU-W1 inverted spinning disk confocal microscope.

For electrophysiology experiments, the penetration depth was estimated *post hoc* using the DiO track of the electrode (see **Electrophysiology**) and the layer 4/layer 5 border was defined based on the DAPI staining. This allowed us to determine which pins of the electrode were located in layer 5/6. For scatter plots, inverse-tuned units (see **Data analysis**) were defined as layer 2/3 or layer 5/6 units if they were above or below this border, respectively.

To identify and quantify inhibitory long-range projections from HVAs to V1 (Extended Data Fig. 7h-l), we injected an AAVretro.CAG.Flex.tdTomato in V1 of GADcre mice and waited approximately 3 weeks before sacrificing the mice. The borders between V1 and the HVAs were defined based on the DAPI staining using the thickness of layer 4. Based on these borders and a mouse atlas^43^, we defined the location and identity of HVAs. To quantify the number of inhibitory neurons in HVAs projecting to V1, we counted the tdTomato-positive cell bodies in the coronal slice that contained the center of the area. Note that this underestimates the difference in the number of projection neurons between V1 and HVAs.

### Data analysis

All data were analyzed using custom-written code in Matlab (MathWorks).

#### Two-photon calcium imaging

We analyzed two-photon calcium imaging data as described previously^44^. Briefly, data were full-frame registered using custom-written software (https://sourceforge.net/projects/iris-scanning/). We selected the neurons semi-manually, based on mean and maximum projection images. We calculated the raw fluorescence traces as the average fluorescence of all pixels within a selected region of interest for each frame. Fluorescence changes (ΔF/F) were calculated as described elsewhere^45^. All stimulus evoked responses were baseline subtracted (1 s pre-stimulus interval).

#### Extracellular recordings

We determined single unit firing using KiloSort and Phy (https://github.com/cortex-lab/KiloSort). We determined the spike times with 1 ms resolution. Inhibitory units were defined as units that significantly increased their firing rate during optogenetic stimulation in the absence of a visual stimulus, i.e. during the pre-visual-stimulus baseline. All stimulus evoked responses were baseline subtracted (0.5 s pre-stimulus interval).

#### Response amplitude

The response amplitude to a stimulus was computed as the average response over the duration of the stimulus presentation (excluding the first 0.5 s of each trial for two-photon experiments due to the delay and slow rise of calcium indicators). Responses were normalized by the maximum response over the relevant stimulus parameter space and then averaged over neurons or units. We defined significant responses as responses that exceeded a z-score of 3.29 (corresponding to p < 10^-3^) or 5.33 (corresponding to p < 10^-7^; for two-photon experiments in layer 4).

#### Receptive field mapping

To estimate the center of the receptive field, we fitted the responses to patches of gratings with a two-dimensional Gaussian. We excluded neurons if they failed to have at least one significant trial-averaged response within 10° of their estimated centers (or the closest data point if no stimulus was located within 10°). For the comparison of the average receptive field maps to classical and inverse stimuli (Fig. 1 and 2, Extended Data Fig. 4 and 5), we only included neurons with at least one significant average response to a classical and an inverse stimulus at any location. To compare regular and fine receptive field mapping (Extended Data Fig. 4), neurons were only included if they responded to both fine and regular grid stimuli and if their estimated receptive field center (of the regular grid) was within the surface covered by the fine mapping stimuli (see smaller dashed rectangle in Extended Data Fig. 4a). To illustrate the average receptive fields (heat maps in Fig. 1, Extended Data Fig. 4 and 5), we used a spline interpolation and smoothed the overall average with a two-dimensional Gaussian filter (10°). We excluded neurons from further analysis (e.g. size tuning, …) if the estimated centers of their ffRFs were not within 10° of the centers of the stimuli presented to establish size tuning, orientation tuning, (…).

#### Size tuning

We fitted the data to an integral over a difference of Gaussians. This fit was used to estimate the neurons’ ffRF and feedback receptive field (fbRF) sizes. We approximated the ffRF size by the size of the patch of gratings evoking the largest response (size tuning fits were bound to the interval 0.1 to 90.1°). Note that we excluded neurons from further analysis if they failed to respond to at least one classical stimulus of any size. To compare size tuning with sharp and burred edges, neurons had to respond to at least one classical stimulus of any size for both stimulus types (sharp and blurred; Extended Data Fig. 1). Surround suppressed neurons were defined as neurons in which the responses to any classical stimulus were significantly larger than those to the largest classical stimulus tested (Extended Data Fig. 5). We calculated the suppression index as the average response over the two largest stimuli presented divided by the maximum response (Extended Data Fig. 8c). The same sizes were used to calculate the suppression index during HVA silencing.

#### Defining inverse-tuned neurons

Neurons were defined as inverse tuned if they significantly responded to at least one classical and one inverse stimulus and if their response to at least one inverse stimulus of any size centered on their ffRF was significantly larger than that to a full-field stimulus (or approximated by the response to the largest classical or smallest inverse stimulus presented).

#### Inverse tuning index (ITI)

We defined the inverse tuning index as:

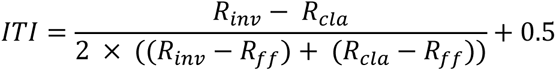

with *R_inv_*: maximum response to inverse stimuli, *R_cla_*: maximum response to classical stimuli, and *R_ff_*: response to a full-field stimulus.

#### Orientation tuning

We fitted a circular sum of Gaussians with a peak offset of 180° and equal tuning width (full width at half maximum of the Gaussian fit). We calculated orientation selectivity index (OSI) and direction selectivity index (DSI) as described elsewhere^12^. Classical and inverse stimuli were presented at a fixed stimulus diameter (10°, 15°, or 20°). Neurons were excluded from this analysis (Extended Data Fig. 3b-g) if their classical and inverse preferred sizes were not within 10° of the presented stimulus size.

#### Contrast tuning

Classical and inverse stimuli were presented at a fixed stimulus diameter (10°, 15°, or 20°) and at one orientation. Neurons were excluded from this analysis (Extended Data Fig. 3h) if their classical and inverse preferred sizes were not within 10° of the presented stimulus size. Moreover, we excluded neurons if their OSIs were above and ≥ 0.3 AND if their orientation preference was not within 45° of the presented stimulus orientation. In other words, we excluded neurons that were strongly orientation tuned to the orthogonal orientation.

#### Response dynamics

To estimate the response delay, rise time, and onset slope for classical and inverse stimuli, we binned the spike times in bins of 10 ms and then median filtered (50 ms) the average traces. We defined the response delay as the first data point after stimulus onset that crossed a z-score threshold of 5.33 (corresponding to p < 10^-7^). Further, we defined the rise time as the interval between the response onset (as estimated for the response delay) and the first time point crossing 75% of the maximum response during stimulus presentation (changing this arbitrary value to 50% or 100% did not affect the results). Finally, we estimated the response onset slope as the fitted slope to the response during the initial rise time. We excluded units whose responses did not exceed the response threshold defined above. Furthermore, for the population responding to the classical stimulus (Fig. 3g), units were excluded if their preferred classical size was larger than the presented stimulus size (±10°). For the inverse-tuned population (Fig. 3g), units were excluded if their preferred inverse size was smaller than the presented size (±10°). For the inverse-tuned subpopulation of units responding to both (Fig. 3c-f), both classical and inverse sizes were required to be within 10° of the presented stimulus size.

#### Awake/anesthetized

Neurons were included in this analysis based on their awake responses (Fig. 4 and Extended Data Fig. 6). However, to ensure that the stimuli were also centered on the receptive fields under anesthesia, neurons were excluded if the estimated centers of their ffRFs under anesthesia were not within 10° of the centers of the anesthetized size tuning stimuli presented. To estimate the peak response of a neuron under anesthesia, we used the same size as in the awake condition (±10°).

#### Size tuning of (non-centered) LM boutons to stimuli centered on their putative V1 targets

Size tuning stimuli were presented at a location such that the population-averaged center of the V1 receptive fields was within 10° (Extended Data Fig. 10f, left). LM boutons were excluded from this analysis if their estimated centers of their ffRFs were within 10° of the centers of the presented size stimuli (Extended Data Fig. 10h). Hence, only putative offset boutons were included. Additionally, for Fig. 5i, boutons needed to respond to an inverse stimulus of any size (stimulus was NOT centered on the boutons’ receptive fields).

#### Intrinsic optical imaging maps

We calculated the temporal phase of the Fourier component at the frequency of the bar presentation. This gave us the complete extent of V1. For locating HVAs, we cross checked the Fourier maps with those obtained from the responses to patches of gratings at different retinotopic locations and confirmed them by standard maps in the literature^46^.

#### Modulation index

We calculated the modulation indexes as the difference between the activity during the optogenetic and the control condition divided by the sum of the two.

### Statistics

We used Wilcoxon rank-sum tests for independent group comparisons, Wilcoxon signed-rank tests for paired tests and Student’s t-tests for a single group analysis. No statistical methods were used to pre-determine sample sizes, but our sample sizes were similar to those used in previous publications.

### Data availability

Datasets supporting the findings of this paper are available on request from the corresponding authors.

### Code availability

Custom code is available from the corresponding authors on request.

---

**Extended Data Figure 1.**
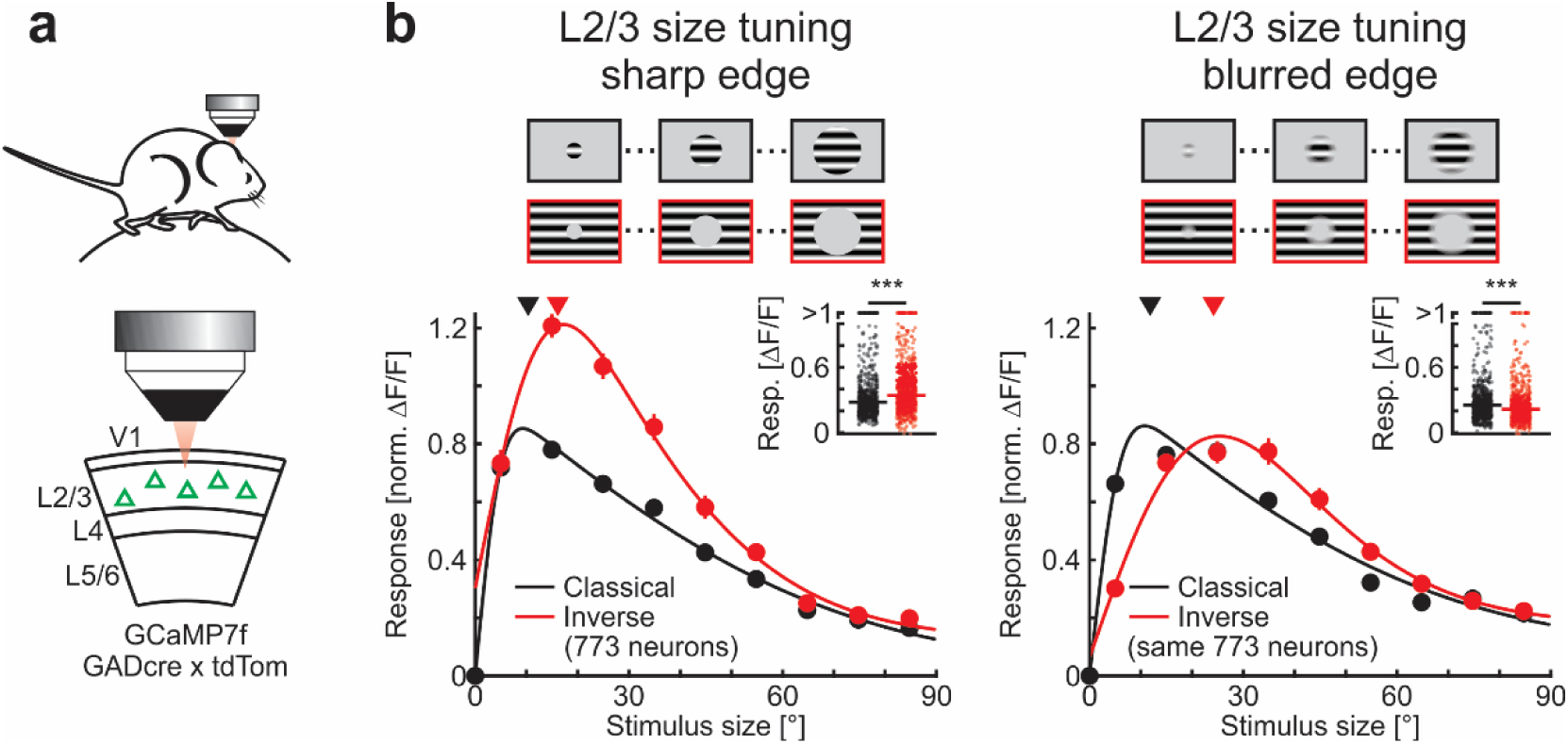
Robust responses to inverse stimuli with blurred edges. **a**, Experimental configuration. **b**, Top: Schematic of stimuli used for size tuning functions. Bottom: Population-averaged size tuning of classical and inverse stimuli with sharp edges (left) and stimuli with blurred edges (right; see Methods). Here and in all other figures, black and red traces are responses to classical and inverse stimuli, respectively, and shaded areas are periods of stimulus presentation. Solid lines are fits to the data (see Methods). Triangles above size tuning functions indicate median preferred size for each condition. Insets: Maximum responses. Horizontal lines, medians. Wilcoxon signed-rank test; sharp edge, ***: p < 10^-8^; blurred edge, ***: p < 10^-9^; 773 neurons in 4 mice. Traces and data points represent mean ± SEM (shading or error bars, respectively). Here and in all other figures, error bars are present but sometimes smaller than symbols.

**Extended Data Figure 2.**
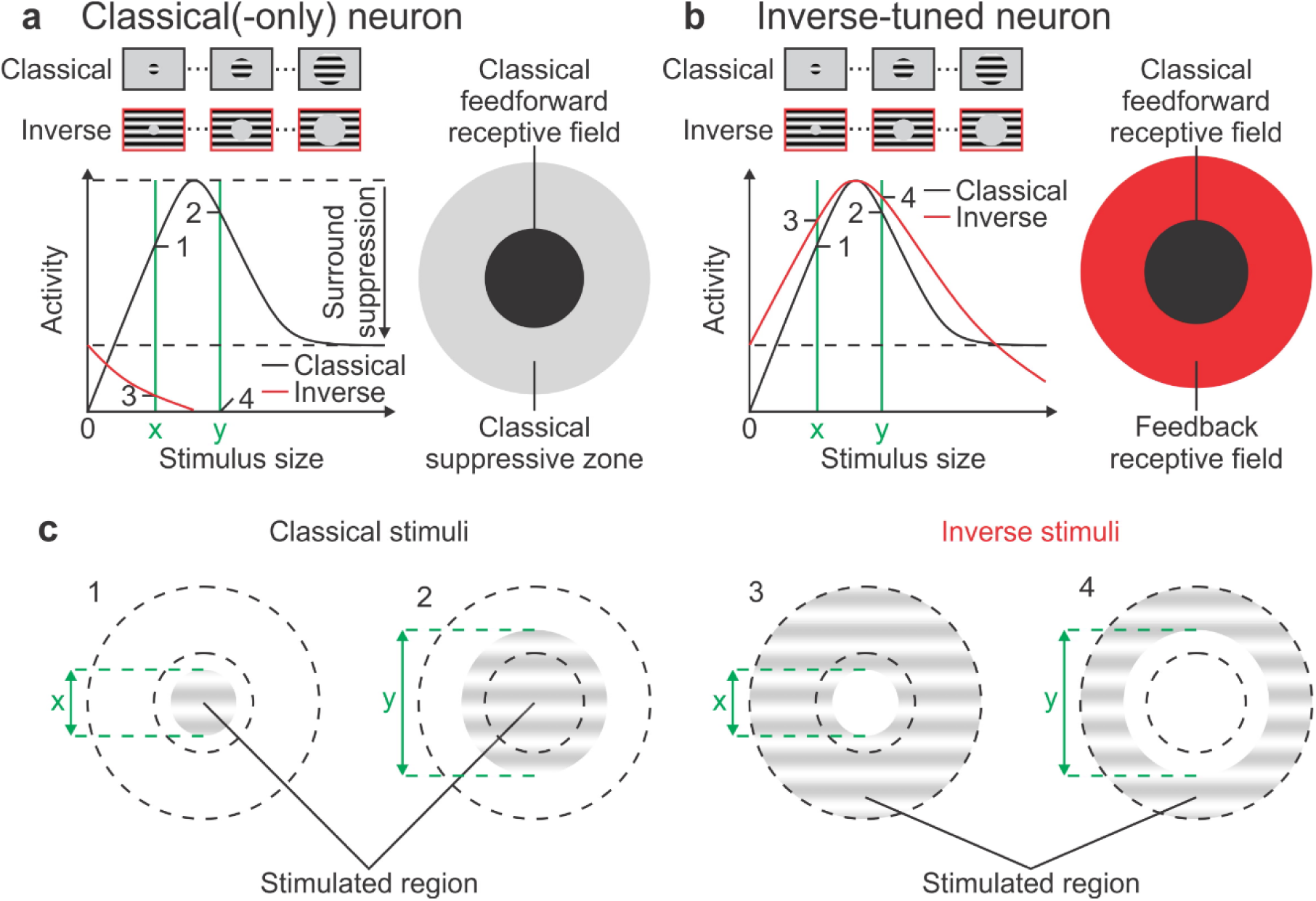
Illustration of classical and inverse-tuned neurons. **a**, Classical(-only) neuron. Left: The response of a neuron probed with classical stimuli (black) increases with the size of the stimulus until it peaks at the neuron’s preferred size (top horizontal dotted line). The response then decreases due to surround suppression (maximum suppressed level indicated by the lower dotted horizontal line). The response of the same neuron probed with inverse stimuli (red) starts at the maximally surround suppressed activity level (an inverse stimulus with a size of 0° is a full-field grating) and then decreases as the diameter of the gray patch increases, consistent with visual stimulation being progressively removed from the classical feedforward receptive field (ffRF). Right: Schematic of a neuron’s ffRF surrounded by its classical suppressive zone**. b**, Inverse-tuned neuron. Left: The response of the neuron probed with inverse stimuli (red) starts, as for the classical-only neuron, at the maximally surround suppressed activity level but then it increases until reaching the neuron’s preferred inverse stimulus size and decreases with larger diameters of the gray patch consistent with visual stimulation being progressively removed from the feedback receptive field (fbRF). Right: Schematic of a neuron’s ffRF surrounded by its fbRF. **c**, Four example stimuli: Two classical stimuli (1 and 2 of sizes x and y, respectively) and two inverse stimuli (3 and 4, also of sizes x and y, respectively). The inner dotted circle represents the outer border of the classical ffRF. The outer dotted circle represents the outer border of the suppressive region and, for inverse-tuned neurons, also the outer border of the fbRF. The response amplitudes to the four example stimuli (1 to 4) in a classical-only neuron and in an inverse-tuned neuron, are marked in (**a**) and (**b**), respectively, at the intersection between the green vertical lines (stimulus size) and the size tuning curves.

**Extended Data Figure 3.**
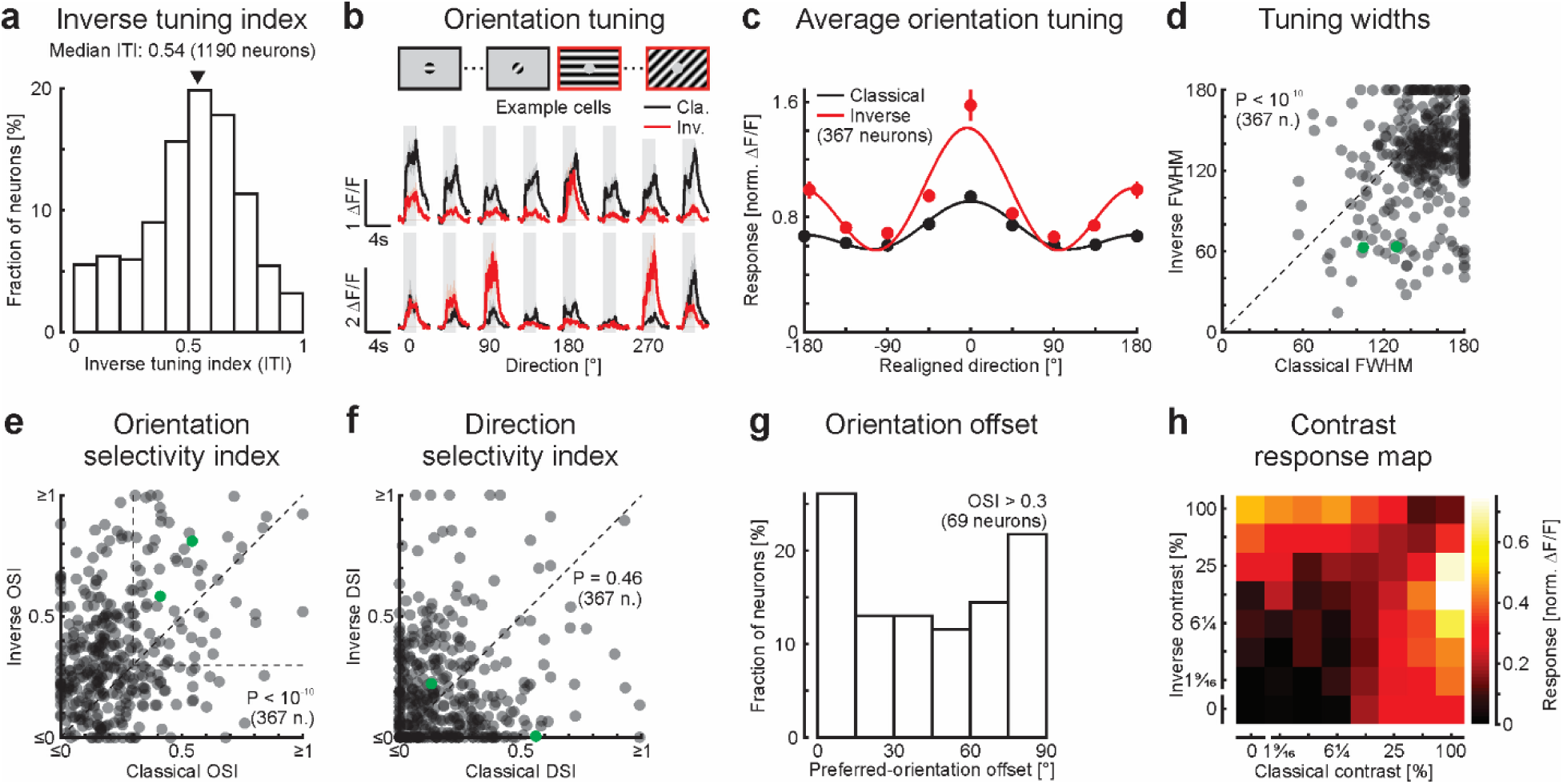
Classical and inverse tuning properties in layer 2/3 excitatory neurons. **a**, Distribution of inverse tuning indices (ITIs) of layer 2/3 excitatory neurons (0: classical only; 0.5 equal peak response to classical and inverse stimuli; 1: inverse only; see Methods). Triangle above the distribution indicates median. Same neurons as in Fig. 1c; 1190 neurons in 9 mice. **b**, Top: Schematic of stimuli presented at different orientations to map the classical and inverse orientation preferences. We tested 8 orientations at intervals of 45° at the neuron’s preferred stimulus size and location using either a classical or an inverse stimulus. Bottom: Calcium responses of two example neurons in primary visual cortex (V1) for each orientation using classical and inverse stimuli. **c**, Population-averaged tuning curve for classical and inverse stimuli. Each neuron’s preferred orientations (independently for classical and inverse stimuli) were aligned to 0° and its activity normalized to its maximum response (367 neurons in 4 mice). Solid lines are fits to the data (see Methods). **d**, Tuning widths of orientation tuning curves obtained with classical stimuli compared to those obtained with inverse stimuli. For each neuron, tuning width was defined as the full width at half maximum (FWHM) of the fitted tuning curve. Wilcoxon signed-rank test; p < 10^-10^; 367 neurons in 4 mice. Green symbols represent the example neurons shown in (**b**). **e**, Same for orientation selectivity indexes. The horizontal and vertical lines at 0.3 delimit the orientation-selective population. Wilcoxon signed-rank test; p < 10^-10^; 367 neurons in 4 mice. **f**, Same for direction selectivity indexes. Wilcoxon signed-rank test; p = 0.46; 367 neurons in 4 mice. **g**, Distribution of orientation offsets. For orientation-selective neurons only (see (**e**), with both OSIs ≥ 0.3), an orientation offset was computed, defined as the absolute difference in orientation between a neuron’s preferred orientation for a classical and an inverse stimulus. **h**, Contrast response map. Classical and inverse stimuli were presented simultaneously, and different combinations of contrasts were tested. The contrast heat map was obtained by averaging normalized activity (86 neurons in 4 mice). Traces and data points represent mean ± SEM (shading or error bars, respectively).

**Extended Data Figure 4.**
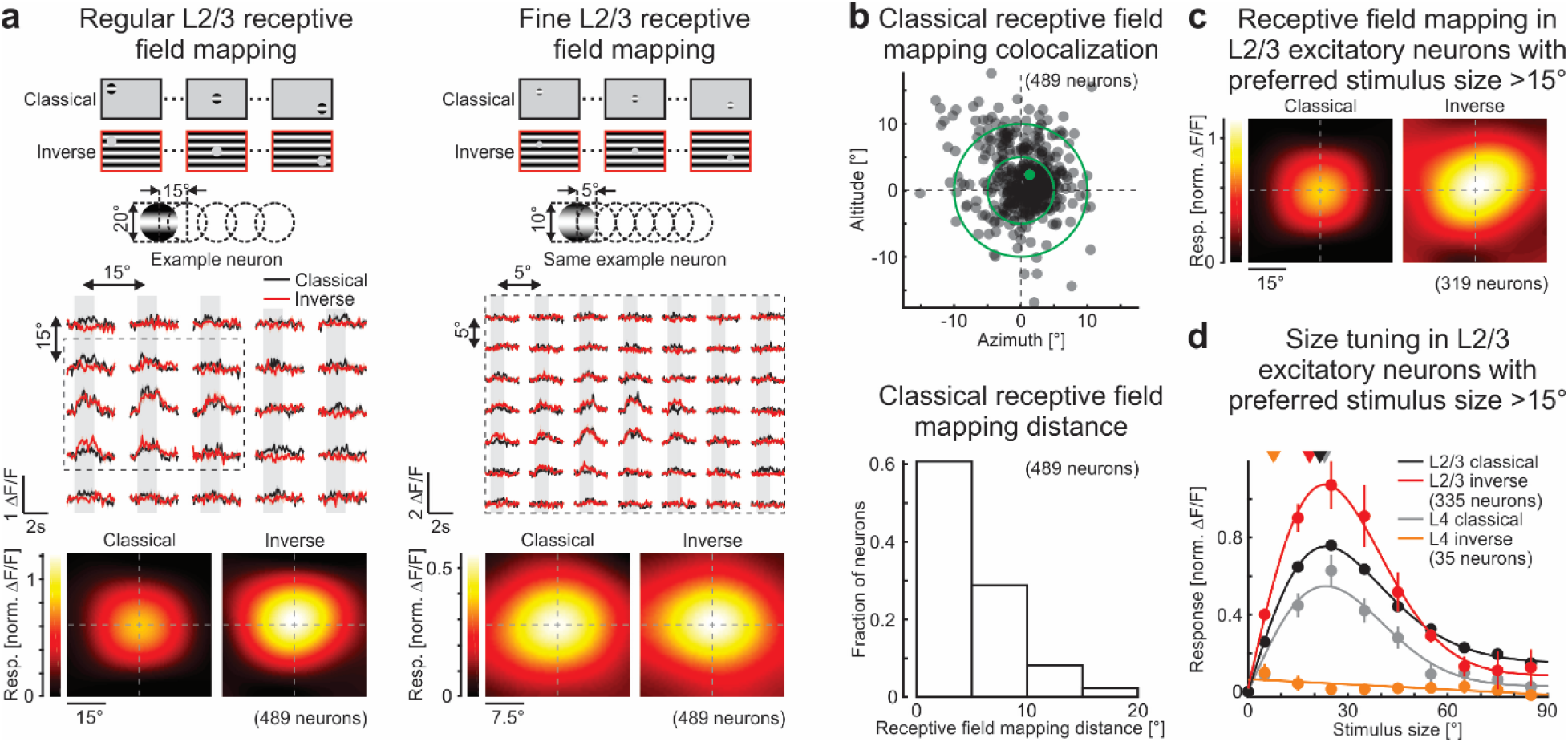
Inverse tuning is not due to low mapping resolution. **a**, Top left: Schematic of regular receptive field mapping. Stimulus diameter of 20° with a grid spacing of 15°. Center left: Trial-averaged calcium responses from an example neuron for each stimulus location. Bottom left: Population-averaged receptive field for responses to classical or inverse stimuli aligned to the center of the ffRF (489 neurons in 4 mice). Right: Same but with fine receptive field mapping. Stimulus diameter of 10° with a grid spacing of 5° (only for part of the visual space covered with the regular mapping, see dotted rectangle on the left). **b**, Top: Spatial offset of regular ffRF mapping compared to fine ffRF mapping (same 489 neurons in 4 mice). For each neuron, its ffRF center estimated by the fine grid mapping is aligned at [0,0] and the localization of its estimated ffRF center estimated by the regular grid is plotted with respect to the fine grid estimated center. Bottom: Distribution of distances between the center of ffRF estimated by fine grid mapping and the center estimated by regular grid mapping (approximately 90% of neurons have a distance between the two centers below 10°). Green symbol represents the example neuron shown in (**a**). **c**, Population-averaged receptive field for responses to classical or inverse stimuli aligned to the center of the ffRF and only for layer 2/3 neurons that had a preferred ffRF size of more than 15° (319 neurons in 9 mice). **d**, Population-averaged size tuning curves for classical (black: layer 2/3 neurons with ffRF > 15°, 335 neurons in 9 mice; gray: layer 4 neurons, 35 neurons in 6 mice) and inverse (red: layer 2/3 neurons with ffRF > 15°, 335 neurons in 9 mice; orange: layer 4 neurons, 35 neurons in 6 mice) stimuli. Solid lines are fits to the data (see Methods). Triangles above size tuning functions indicate median preferred size for each condition. Inset: Maximum responses. Horizontal lines, medians. Layer 2/3 neurons with ffRF size > 15°; Wilcoxon signed-rank test; *: p = 6.7 × 10^-3^; 335 neurons in 9 mice. Traces and data points represent mean ± SEM (shading or error bars, respectively).

**Extended Data Figure 5.**
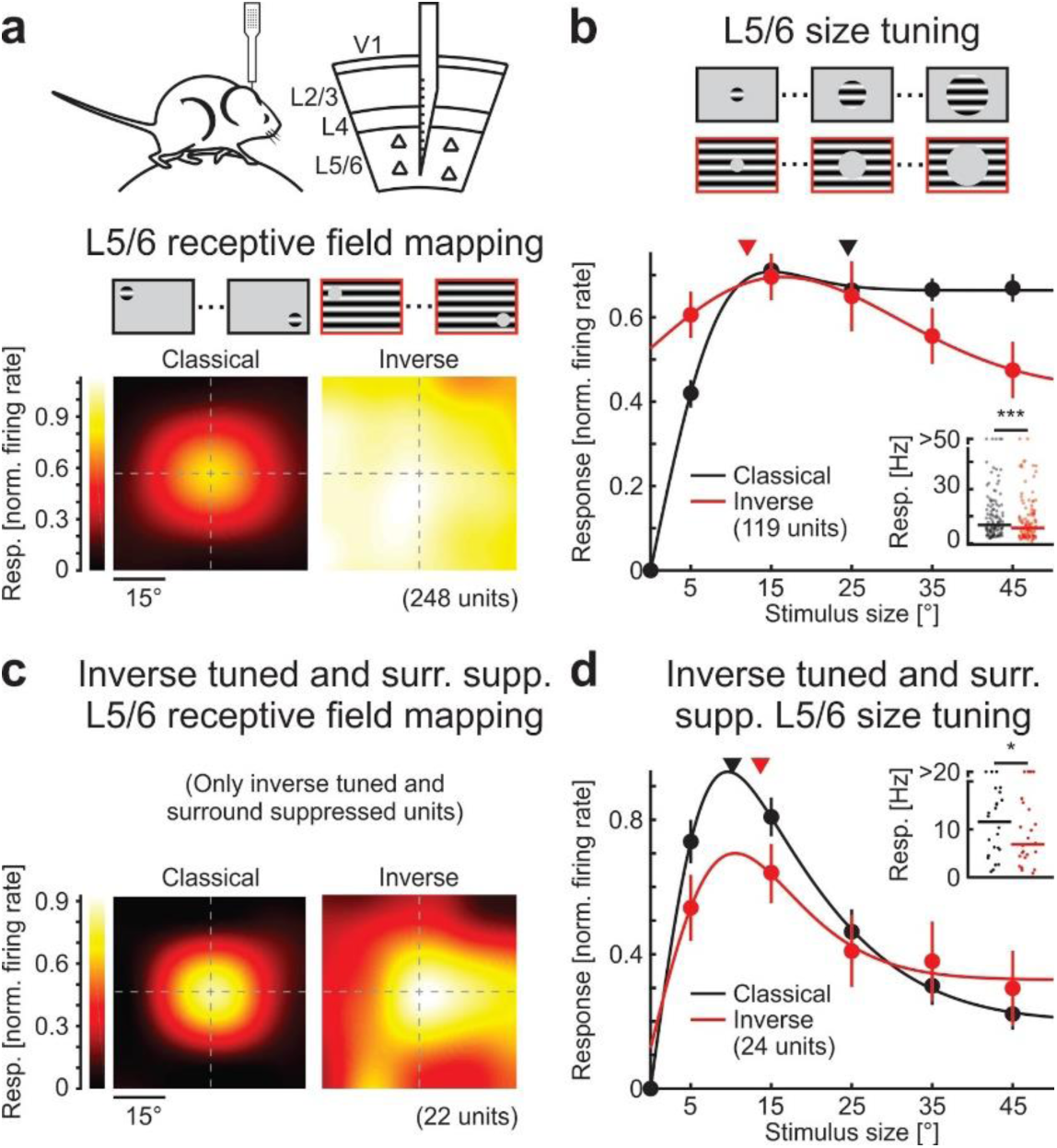
Responses to inverse stimuli in layer 5/6. **a**, Receptive field mapping of layer 5/6 neurons using classical and inverse stimuli. Top: Experimental configuration. Electrophysiological recordings were obtained in awake mice. The silicon probe spanned all layers, including deep layers (see Methods for layer definition). Center: Receptive fields were mapped using classical and inverse stimuli. Bottom left: Population-averaged ffRFs for layer 5/6 neurons. Bottom right: Same for inverse stimuli, aligned relative to the center of the ffRF (248 units in 20 mice). **b**, Size tuning of layer 5/6 neurons using classical and inverse stimuli. Top: Schematic of stimuli used for size tuning functions. The classical and inverse stimuli were presented at the same location (within 10° of the estimated center of the ffRF). Bottom, normalized size tuning curves for classical and inverse stimuli. Solid lines are fits to the data (see Methods). Triangles above size tuning functions indicate median preferred size for each condition. Inset: Maximum responses. Horizontal lines, medians. Wilcoxon signed-rank test; ***: p = 1.1 × 10^-4^; 119 units in 20 mice. **c** and **d**, same as (**a** and **b**) but for subset of layer 5/6 units defined both as surround suppressed and inverse tuned (as compared to (**b**) where all layer 5/6 units that responded to at least one classical stimulus size were included; see Methods) (**c**), 22 units in 12 mice; (**d**), Wilcoxon signed-rank test; *: p = 0.016; 24 units in 12 mice. Data points represent mean ± SEM (error bars).

**Extended Data Figure 6.**
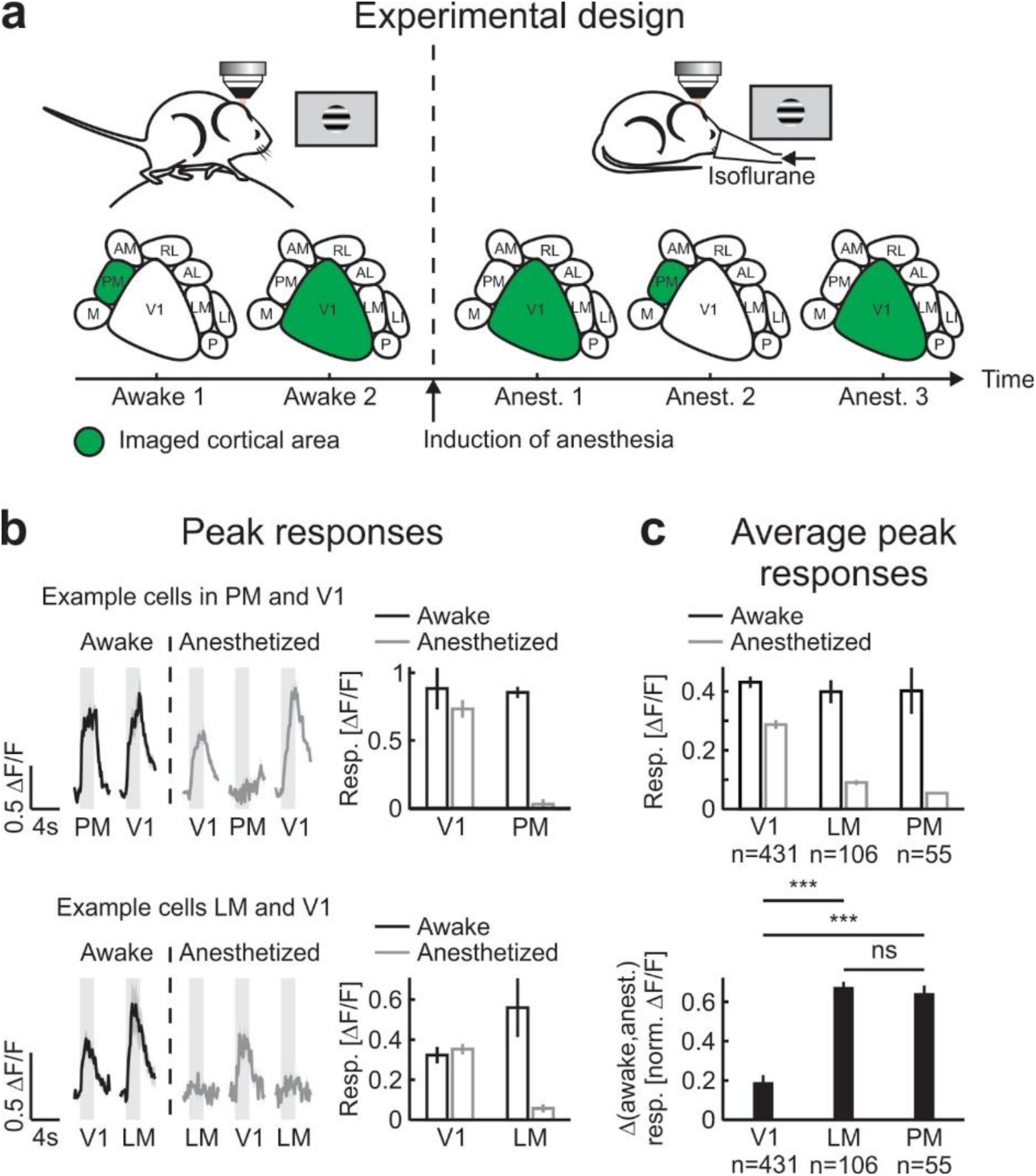
Impact of anesthesia is more pronounced in higher visual areas. **a**, Experimental design. The responses to classical stimuli of neurons in a higher visual area (HVA), LM or PM, and V1 were recorded using two-photon calcium imaging. The experiment started in awake mice by imaging either a HVA or V1. After induction of anesthesia, the same neurons were imaged again. To reduce the influence of variability in anesthesia levels, the first imaged area under anesthesia was imaged again at the end of the experiment. **b**, Peak responses in visual areas. Top left: Example calcium response of a neuron located in PM and another neuron located in V1 in an awake mouse (black) and responses of the same neurons in the anesthetized mouse (gray). Top right: Trial-averaged peak response for the same neurons shown on the left for the awake (black) and anesthetized mouse (gray). Bottom, same for a different mouse but recorded in V1 and LM. **c**, Population-averaged peak responses in awake and anesthetized mice. Top: Population-averaged peak responses in V1, LM and PM for awake (black) and anesthetized mice (gray). Wilcoxon signed-rank test; V1, p < 10^-10^, 431 neurons in 5 mice; LM, p < 10^-10^, 106 neurons in 3 mice; PM, p < 10^-9^; 55 neurons in 2 mice. Bottom: Population-averaged difference between normalized neuronal activity for awake and anesthetized state. For each neuron, all responses were normalized by the peak activity in the awake state before computing the differences. Wilcoxon rank-sum test; V1, 431 neurons in 5 mice; LM, 106 neurons in 3 mice; PM, 55 neurons in 2 mice. V1-LM, ***: p < 10^-10^; V1-PM, ***: p < 10^-10^; LM-PM, NS: p = 0.48. Traces and bars represent mean ± SEM (shading or error bars, respectively).

**Extended Data Figure 7.**
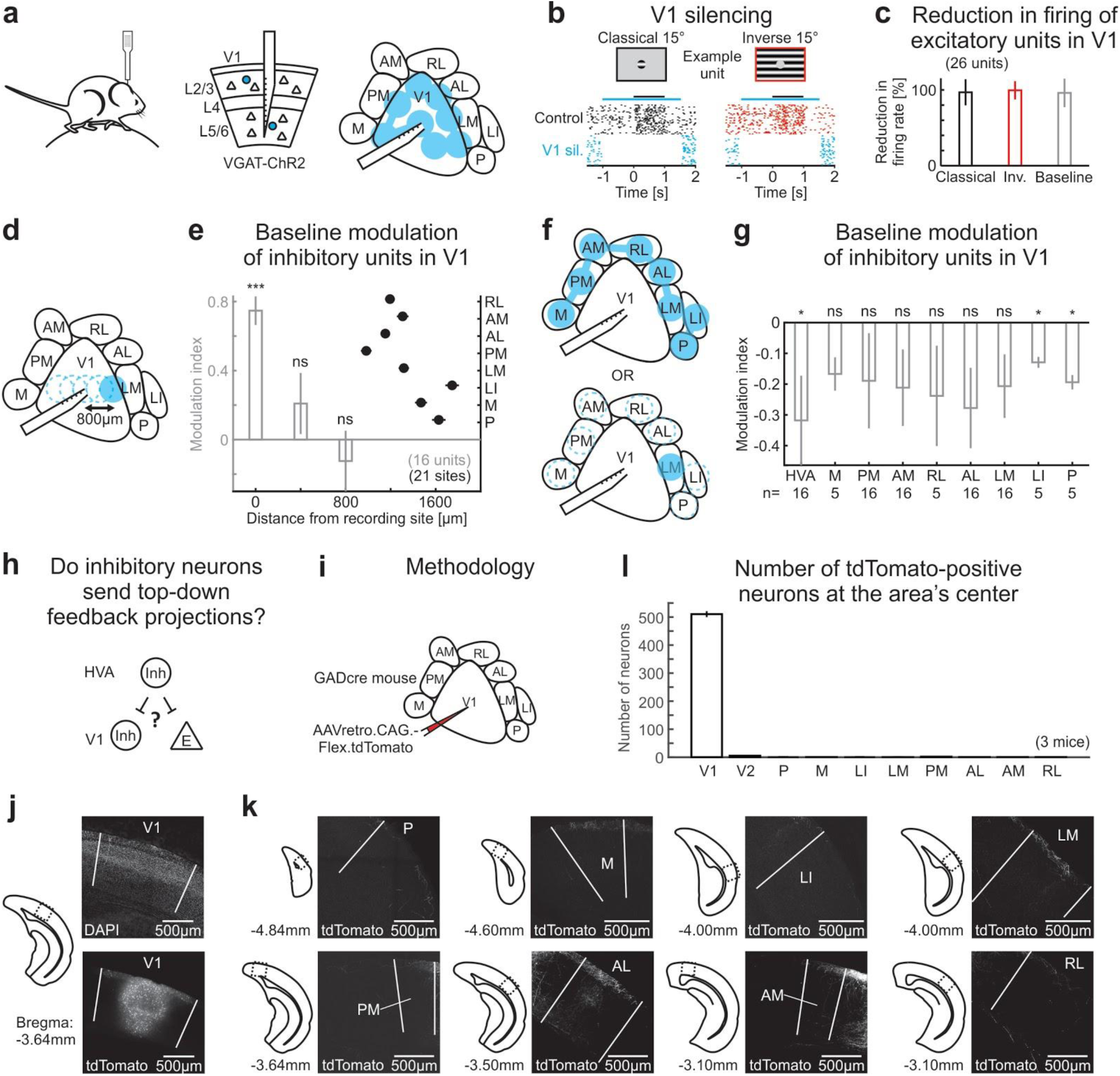
Strong silencing by spatially restricted excitation of local inhibitory units. **a**, Experimental configuration. A silicon probe was inserted in V1, spanning all cortical layers, in mice expressing Channelrhodopsin-2 in inhibitory neurons (VGAT-ChR2). To assess the strength of inhibition of excitatory units when using the laser scanning technique (Fig. 5; see Methods), the V1 recording site as well as seven other locations were scanned at 125 Hz. **b**, Raster plot of example excitatory unit in layer 5/6 in response to classical and inverse stimuli of 15° in diameter under control conditions (30 trials each) and during silencing of V1 (blue; V1 sil.). Black and blue horizontal lines are periods of stimulus presentation and V1 silencing, respectively. Classical and inverse stimuli were presented in random order; trials with V1 silencing were randomized as well but are separated here for clarity. **c**, Reduction in firing of excitatory units. The reduction in firing was measured as one minus the ratio between the optogenetic condition and the control condition. Note that silencing reached nearly 100% for both responses to classical and inverse stimuli, and for the baseline activity (26 units in 10 mice). **d**, Experimental configuration. To assess the effect of distance on the optogenetic stimulation of inhibitory units at the recording site, two medial and two lateral locations at 400 μm and 800 μm from the V1 recording site were targeted for laser stimulation while recording in V1. **e**, Modulation of the baseline of inhibitory units. The modulation index was defined as the difference between the activity during the optogenetic and the control condition divided by the sum of the two. The modulation index was high at the recording site (at 0 μm) and quickly dropped with distance (gray bars; Student’s t-test; 0 μm, ***: p < 10^-6^; 400 μm, ns: p = 0.26; 800 μm, ns: p = 0.51; 16 units in 8 mice). As a comparison, the distance of the HVAs from the recording site is plotted on the same axis (black dots, right y-axis; 21 recording sites, 12 mice), suggesting that when pointing the laser at HVAs, direct activation of inhibitory neurons at the V1 recording site is unlikely. **f**, Experimental configurations. To assess the effect of the laser stimulation of HVAs on inhibitory units at the recording site, all 8 (top) or individual HVAs (bottom) were targeted for laser stimulation while recording in V1 (same configurations as during the experiments in Fig. 5 and Ext. Data Figs. 8 and 9). **g**, Modulation of the baseline of inhibitory units. The modulation indices were either negative or not significantly different from zero, indicating that the laser stimulation was unlikely to directly activate inhibitory neurons at the V1 recording site. Student’s t-test; HVA, *: p = 0.045; 16 units in 8 mice; M, ns: p = 0.16; 5 units in 4 mice; PM, ns: p = 0.24; 16 units in 8 mice; AM, ns: p = 0.11; 16 units in 8 mice; RL, ns: p = 0.46; 5 units in 4 mice; AL, ns: p = 0.051; 16 units in 8 mice; LM, ns: p = 0.064; 16 units in 8 mice; LI, *: p = 0.015; 5 units in 4 mice; P, *: p = 0.010; 5 units in 4 mice. **h**, Are there many inhibitory neurons projecting from HVAs to V1? **i,** Methodology. A retrograde virus, AAVretro.CAG.Flex.tdTomato, was injected in V1 of GADcre mice to label GAD-positive neurons projecting to the site of injection. **j**, Left: Outlines of the cortical section where the confocal images shown on the right were acquired. The location of the imaged area is further indicated by the dotted square depicted on the outline. Rostro-caudal distance to bregma is indicated below the outline. Right: Average intensity projection. Top right: DAPI staining highlights the higher density of neurons in layer 4 in V1 used to define V1 borders (white lines). Bottom: The tdTomato fluorescence reveals numerous cell bodies in V1 around the site of injection and even more distal in L1. **k**, Same as in (**j**) but only for the tdTomato fluorescence and for all HVAs targeted for laser stimulation in Fig. 5 and Ext. Data Figs. 8 and 9. White lines delimit area’s boundaries. **l**, Quantification of tdTomato positive neurons at the area’s center. The number of tdTomato positive neurons were counted in the section containing the center of the investigated area. Note the sparse inhibitory projections from HVAs to V1 but the abundance of local inhibitory projections within V1 (3 mice). Bars represent mean ± SEM (error bars).

**Extended Data Figure 8.**
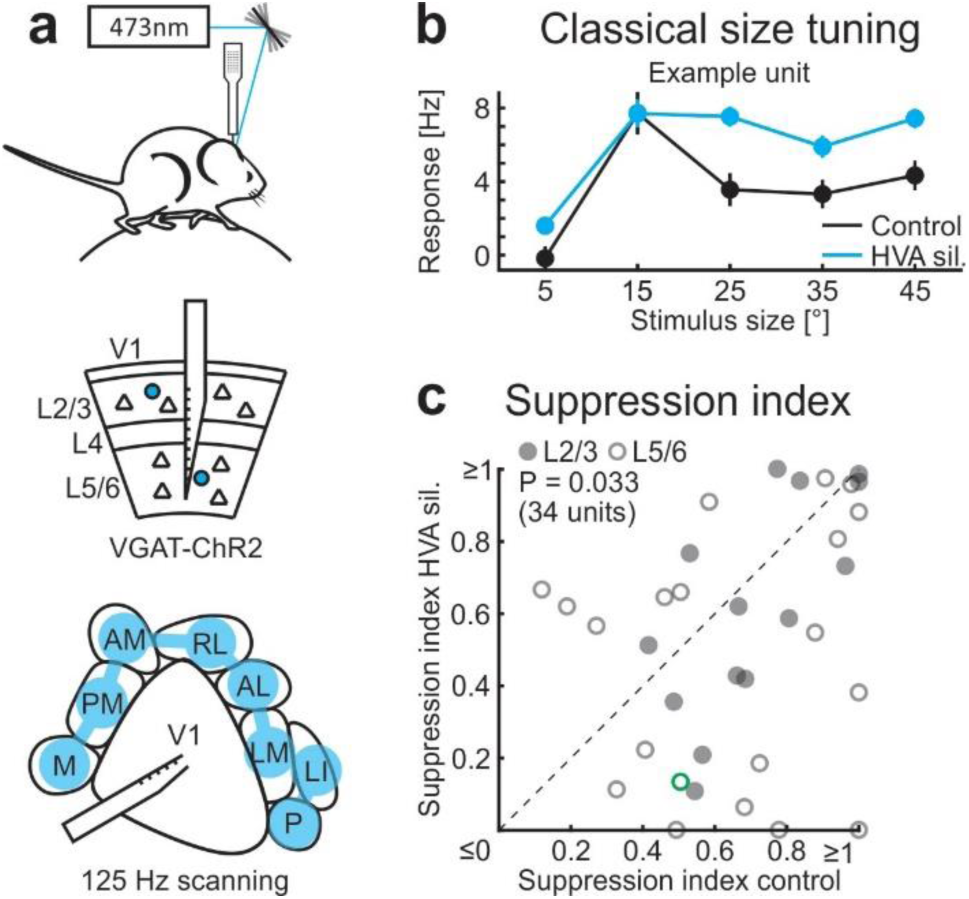
Silencing higher visual areas reduces surround suppression in V1. **a**, Experimental configuration. A laser beam is scanned over HVAs around V1 for optogenetic silencing while recording in V1. **b**, Size tuning function of an example unit (baseline subtracted firing rates) to classical stimuli with (blue) or without (black) HVA silencing. Note the relief of surround suppression at larger stimulus sizes upon silencing HVAs. **c**, Scatter plot of the classical suppression index with or without silencing of HVAs (see Methods). Wilcoxon signed-rank test; p = 0.033; 34 units in 12 mice. Closed and open symbols are units from layer 2/3 and 5/6, respectively. Green symbol represents the example neuron shown in (**b**). Data points represent mean ± SEM.

**Extended Data Figure 9.**
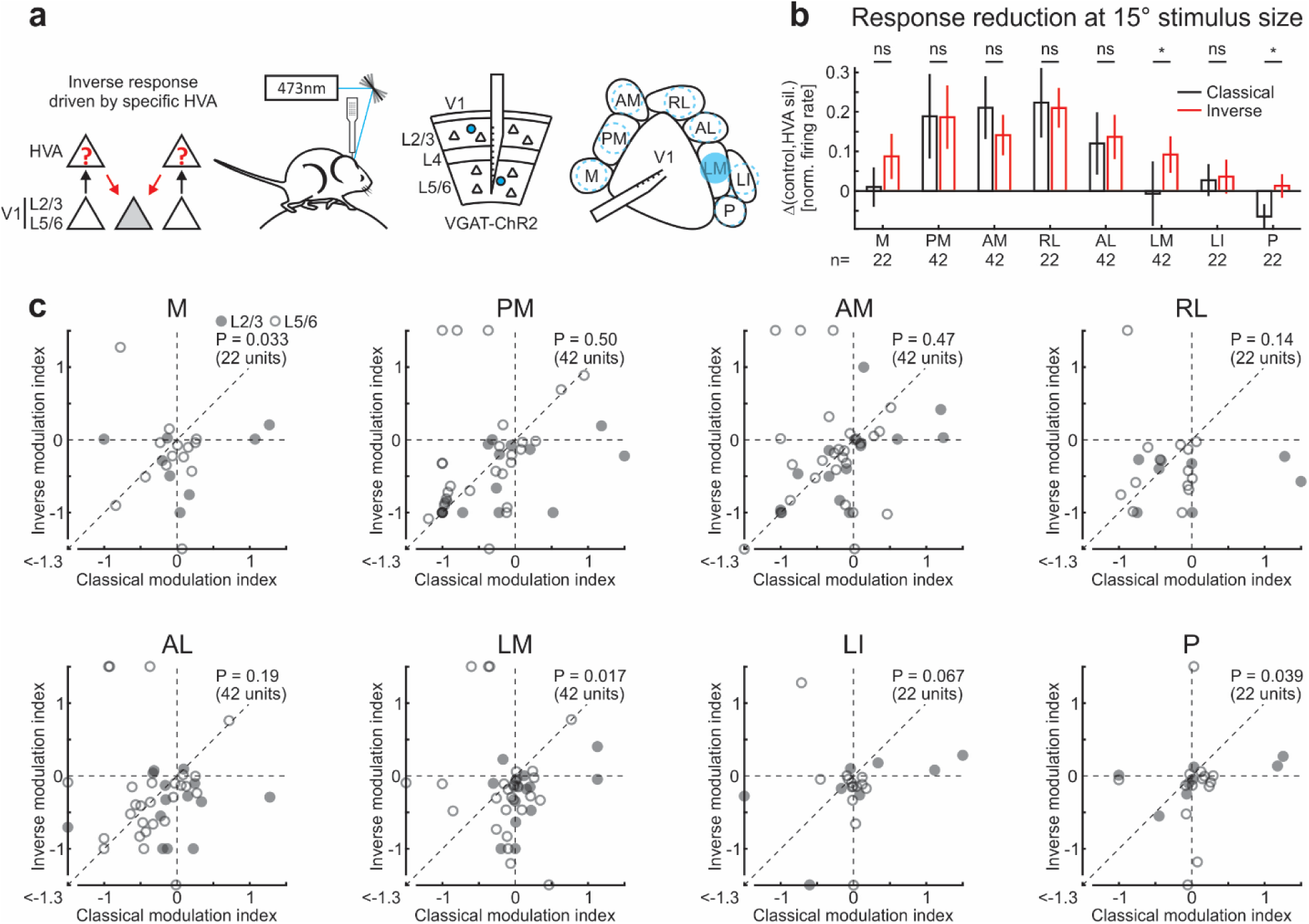
Silencing individual higher visual areas differentially affects responses to classical and inverse stimuli. **a**, Schematic of results and experimental configuration. Individual HVAs are targeted for optogenetic silencing while recording in V1. **b**, Difference in firing rates (baseline subtracted and normalized) between control conditions and individual HVA silencing for classical and inverse stimuli. Wilcoxon signed-rank test; M, ns: p = 0.18; 22 units in 5 mice; PM, ns: p = 0.46; 42 units in 12 mice; AM, ns: p = 0.88; 42 units in 12 mice; RL, ns: p = 0.81; 22 units in 5 mice; AL, ns: p = 0.20; 42 units in 12 mice; LM, *: p = 0.013; 42 units in 12 mice; LI, ns: p = 0.51; 22 units in 5 mice; P, *: p = 0.020; 22 units in 5 mice. **c**, Scatter plot of the modulation indexes of individual HVA silencing for responses to classical and inverse stimuli (see Methods). Closed and open symbols are units from layer 2/3 and 5/6, respectively. Wilcoxon signed-rank test; M, p = 0.033; 22 units in 5 mice; PM, p = 0.50; 42 units in 12 mice; AM, p = 0.47; 42 units in 12 mice; RL, p = 0.14; 22 units in 5 mice; AL, p = 0.19; 42 units in 12 mice; LM, p = 0.017; 42 units in 12 mice; LI, p = 0.067; 22 units in 5 mice; P, p = 0.039; 22 units in 5 mice. Note that for the visual stimulus parameters used here, LM showed the strongest effect in preferentially reducing responses to inverse stimuli. Bars represent mean ± SEM (error bars).

**Extended Data Figure 10.**
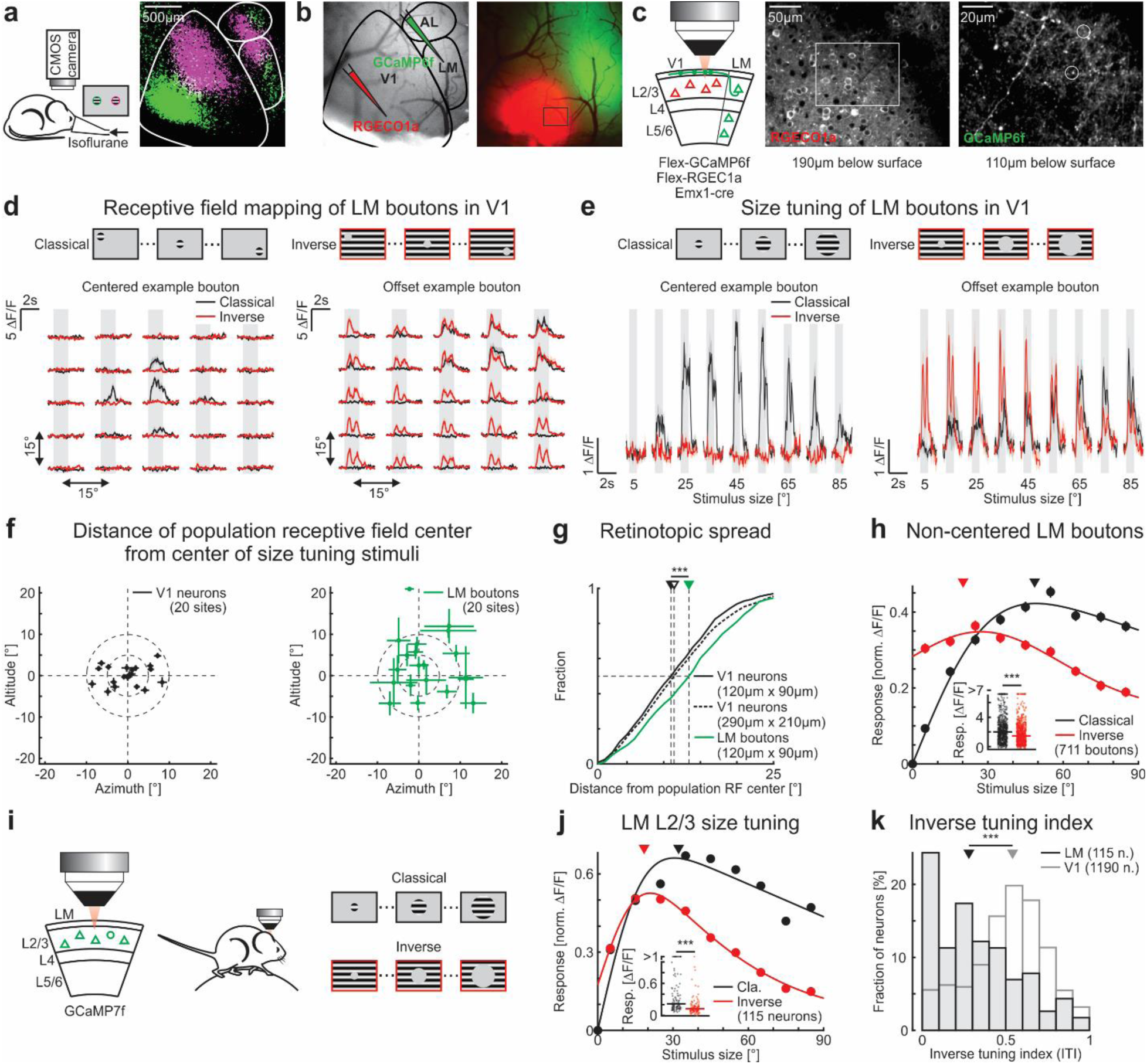
Dual-color imaging of LM boutons and their putative V1 targets. **a**, Left: Experimental configuration. To localize V1 and LM, we used intrinsic optical imaging (see Methods). Right: Response map to a nasal (magenta) and temporal patch of gratings (green). White lines represent area borders. **b**, Left: Blood vessel pattern overlaid with area borders defined by the intrinsic map (black lines). The red-shifted calcium indicator RGECO1a was injected in V1 and GCaMP6f was injected in LM. Right: Fluorescence of calcium indicators in V1 and LM. The black square delimits the example imaging site shown in (**c**). Same scale bar as in (**a**). **c**, Left: The responses of LM boutons and of V1 cell bodies were recorded within the same cortical location. Center: Example imaging site of V1 cell bodies recorded 190 μm below surface. The white square delimits the example imaging site shown on the right. Right: Example imaging site of LM boutons in V1 recorded 110 μm below surface. **d**, Top: Schematic of receptive field mapping. Left: Trial-averaged calcium responses from an example LM bouton aligned to its putative V1 target. Right: same but from an example bouton that is retinotopically offset with respect to its putative V1 target. **e**, Top: Schematic of stimuli used for size tuning functions. Left and right: Trial-averaged calcium responses from the same example neurons as in (**d**). **f**, Left: Distance of population-averaged receptive field center of V1 neurons from center of size tuning stimuli (20 sites in 5 mice). Right: same for LM boutons. Note that all average V1 receptive field centers are located within 10° and that average LM receptive field centers are more spread with larger standard deviations. **g**, Retinotopic spread measured as cumulative distance from population-averaged receptive field center. The ffRF centers of LM boutons (solid green line) were more retinotopically spread than V1 neurons measured over the same cortical surface (solid black line) or measured over approximately 6 times the surface of the LM bouton site (dotted black line). Wilcoxon rank-sum test; LM-V1 same surface, ***: p < 10^-4^; LM-V1 6× surface, ***: p = 3.1 × 10^-4^; LM, 311 boutons in 5 mice; V1 same surface, 530 neurons in 5 mice; V1 6× surface, 2352 neurons in 5 mice. **h**, Population-averaged size tuning function of LM boutons (711 boutons in 5 mice) that are NOT retinotopically aligned with their V1 target. Note that both classical and inverse stimuli were presented at the ffRF location of their putative V1 targets (see Methods) and NOT at the ffRF location of the imaged LM boutons. Solid lines are fits to the data (see Methods). Triangles are median preferred size. Insets: Maximum responses. Horizontal lines, medians. **i**, Experimental configuration. **j**, Population-averaged size tuning functions for classical and inverse stimuli. Solid lines are fits to the data (see Methods). Triangles are median preferred size. Insets: Maximum responses. Horizontal lines, medians. Wilcoxon signed-rank test; ***: p < 10^-9^; 115 neurons in 3 mice. Data points represent mean ± SEM (error bars, respectively). **k**, Distribution of inverse tuning indices (ITIs) of LM (black) and V1 neurons (gray; same neurons as in Fig. 1c; 0: classical only; 0.5 equal peak response to classical and inverse stimuli; 1: inverse only; see Methods). Triangles above the distribution indicate medians. Wilcoxon rank-sum test; ***: p < 10^-10^; 115 neurons in 3 mice and 1190 neurons in 9 mice for LM and V1, respectively.

